# Fine-scale contemporary recombination variation and its fitness consequences in adaptively diverging stickleback fish

**DOI:** 10.1101/2023.07.22.550174

**Authors:** Vrinda Venu, Enni Harjunmaa, Andreea Dreau, Shannon Brady, Devin Absher, David Kingsley, Felicity Jones

## Abstract

Despite deep evolutionary conservation, recombination varies greatly across the genome, among individuals, sexes, and populations and can be a major evolutionary force in the wild. Yet this variation in recombination and its impact on adaptively diverging populations is not well understood. To elucidate the nature and potential consequences of recombination rate variation, we characterized fine-scale recombination landscapes by combining pedigrees, functional genomics and field fitness measurements in an adaptively divergent pair of marine and freshwater threespine stickleback populations from River Tyne, Scotland. Through whole-genome sequencing of large nuclear families, we identified the genomic location of almost 50,000 crossovers and built recombination maps for 36 marine, freshwater, and hybrid individuals at 3.8 kilobase resolution. Using these maps, we quantified the factors driving variation in recombination rate: we find strong heterochiasmy between sexes (68% of the variation) but also differences among ecotypes (21.8%). Hybrids show evidence of significant recombination suppression, both in overall map length and in individual loci. We further tested and found reduced recombination rates both within single marine–freshwater adaptive loci and between loci on the same chromosome, suggestive of selection on linked ‗cassettes‘. We tested theory supporting the evolution of linked selection using temporal sampling along a natural hybrid zone, and found that recombinants with shuffled alleles across loci show traits associated with reduced fitness. Our results support predictions that divergence in cis-acting recombination modifiers whose mechanisms are disrupted in hybrids, may have an important role to play in the maintenance of differences among adaptively diverging populations.

## Introduction

Meiotic recombination is a key source of genetic diversity that shapes the genomic landscape and the evolutionary process. By shuffling parental alleles to produce novel haplotypes, recombination impacts the strength of selection acting on nearby polymorphisms, and can increase the efficacy of selection and adaptation in natural populations. In most sexually reproducing organisms, recombination is essential for proper chromosomal segregation during meiosis. Defects in recombination can have serious phenotypic consequences: inviable gametes, miscarriages, birth defects and developmental abnormalities (1, 2). For these reasons, recombination is thought to be highly constrained (3) — an idea supported by the deep conservation of proteins involved in homologous recombination across all domains of life.

Despite these constraints, recombination varies greatly: significant and heritable differences in recombination location and rate have been observed among individuals (4, 5), sexes (6–9), and species (10–12). Across the genome, recombination rate can vary by orders of magnitude (13, 14) - a phenomenon in some species that involves ‗hot-‘ and ‗coldspots‘ (4, 15–17). Understanding the determinants of this non-random spatial distribution has been one of the core objectives of recombination studies. Evidence suggests that a hierarchical combination of cis-acting factors including chromatin state, epigenetic factors, and DNA sequence context are predictive of crossover location and frequency (18–21). In addition, a number of trans-acting recombination modifiers influencing genome-wide recombination rate (e.g. RNF212 (22–24)) or local genomic recombination landscapes (e.g. PRDM9 (25–27)) have been identified.

Recombination rate variation at different levels causes uneven shuffling of genomic regions, which inevitably shapes the distribution of genetic variation visible to natural selection, and can place recombination modifiers themselves under selection (28–30). Next generation sequencing data has revealed that adaptation in natural populations involves multiple loci across the genome (31) and these loci are often clustered (32–35). This dispersed but non-random distribution implies uneven linkage between adaptive loci, and raises further questions: if recombination is an essential process in meiosis, what are the molecular and functional constraints? What are the effects of shuffling alleles among adaptive loci? How do selective forces shape recombination variation?

Threespine stickleback fish offer an attractive system with several major advantages to investigate how molecular constraints and evolutionary forces interact to shape recombination in natural populations. First, there are a number of stickleback marine–freshwater hybrid zones, where both the number and location of hundreds of divergently adaptive alleles are well-characterized at kilobase resolution (32, 36). Second, we can readily raise field and laboratory crosses with large clutch sizes to quantify individual recombination maps. Direct detection of crossover events from pedigrees is not confounded by population demography and selection, thereby providing an accurate snapshot of the contemporary recombination landscape and complementing widely used LD estimates of recombination in natural populations. Third, there is now a mature genomic and molecular toolkit, including a high-quality, well-annotated genome assembly with variation and functional data to help us interpret our results. Finally, we have access to field sites that allow us to track and monitor allele frequencies over the years, especially in hybrid fish, in order to assess fitness-related traits in nature.

Following the latest glacial retreat from the Northern Hemisphere (approximately 10-20 thousand years ago) ancestral marine sticklebacks colonized and adapted to newly formed freshwater lakes and rivers through parallel changes in numerous morphological, physiological, and behavioral traits (37). This rapid adaptive divergence draws on standing genetic variation at more than 242 loci across the genome (FDR 0.05; (32)) via ongoing gene exchange through hybrid zones (38, 39). In such diverging populations that experience high levels of migration and gene exchange, natural selection may theoretically favor the maintenance of linkage among alleles within ’adaptive cassettes’ (40, 41). If the continued gene flow is not only homogenizing divergence but also bringing beneficial alleles into the population, a recombination modifier with an underdominant mode of action (where recombination suppression occurs only in hybrids) may be particularly favored. Further, in a metapopulation exposed to heterogeneous selection regimes, increased variation in recombination rate itself may be beneficial, e.g. via differences between sexes (heterochiasmy),— variable recombination rates might help populations maintain adaptation to their local environment while navigating the costs and benefits of gene flow involving both beneficial and deleterious alleles from diverged neighboring populations. In sticklebacks, several studies have confirmed strong heterochiasmy and that divergently adaptive loci fall into regions of low recombination (34, 42, 43) but it remains unclear whether recombination rates have become locally modified or whether molecular constraints on recombination rates have dictated the locations where adaptive cassettes can form.

In species lacking PRDM9, local recombination rates have been assumed to be highly conserved (44). However, although sticklebacks lack a functional copy of PRDM9, their recombination hotspots have been shown to vary considerably between adaptively diverged populations (45). Neither the fitness consequences nor the molecular mechanisms of this variation have been addressed at high resolution. In this study we contrast high-resolution genomic recombination maps of marine, freshwater and hybrid ecotypes with genomic and molecular features associated with crossovers, and explore the potential fitness implications of recombination among linked adaptive loci in admixed individuals from a hybrid zone. Combined, our results provide a unique platform for exploring the interplay of recombination and natural selection in adaptively diverging populations.

## Results

### Strong female-biased heterochiasmy in individual maps of recombination

We built high-resolution, genome-wide maps of recombination crossover locations for each of 36 individuals from wild-derived strains of marine, freshwater, and F1 hybrid fish using whole-genome sequencing followed by whole-chromosome haplotype phasing of large nuclear families with 90-94 offspring (Fig 1a-c). Offspring of a nuclear family carry recombinant chromosomes generated during maternal and paternal gametogenesis, and collectively harbor thousands of recombination crossovers from each parent (>2800 per family). From 3,338 meiotic products, we assembled a comprehensive set of 49,838 meiotic crossover locations (18,039 paternal and 31,809 maternal) with median resolution of 3845 bp (Fig S1). The observed number of crossovers corresponds to a mean sex-averaged genetic map length of 1493 cM and a genome-wide average recombination rate of 3.24 cM/Mb, broadly consistent with map lengths in previous studies (1404 cM, range: 993– 1963cM, from (9, 42, 46–48).

**Fig 1:**
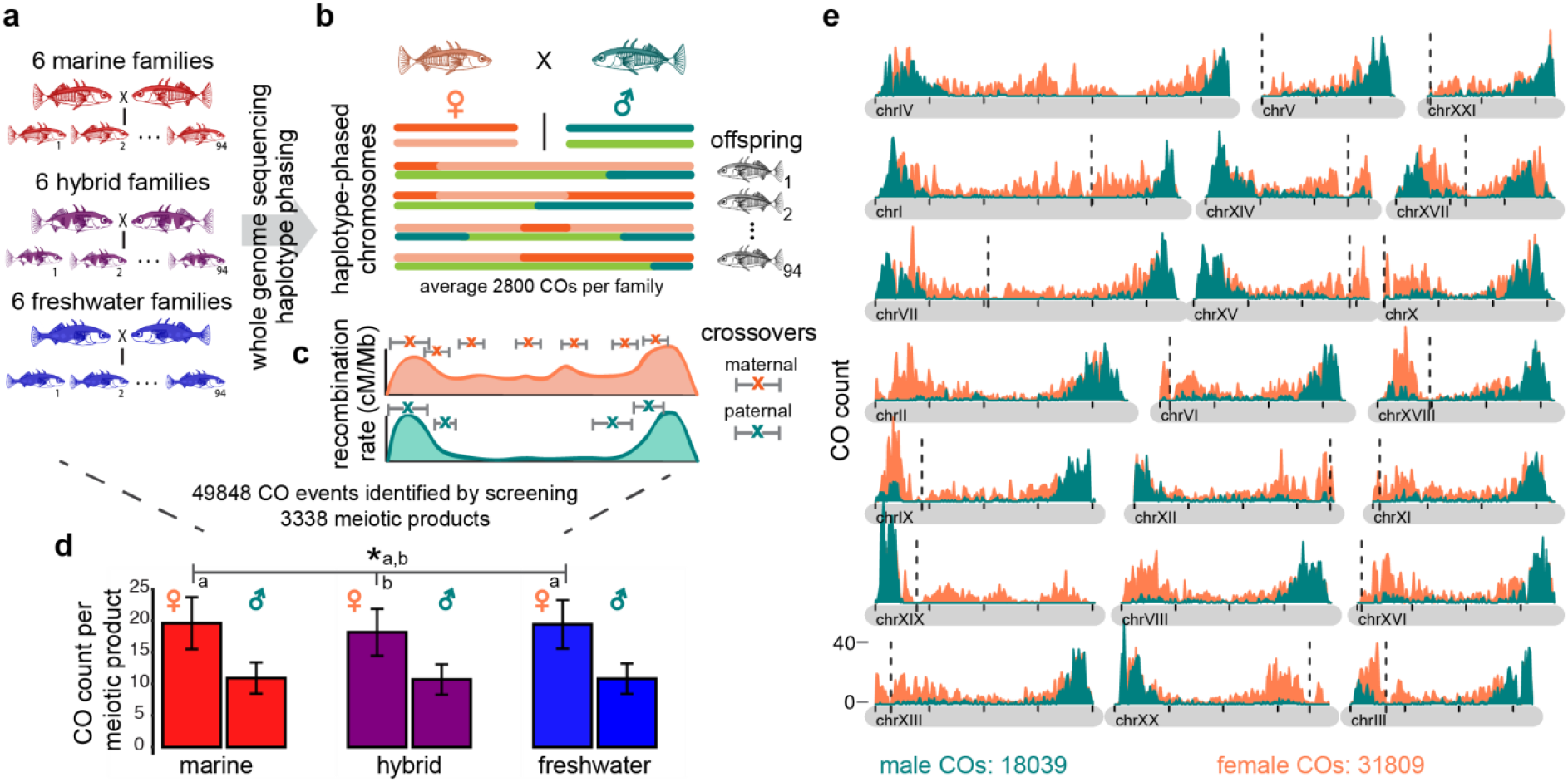
Construction of single-individual recombination maps by nuclear family whole genome sequencing. a) 18 large clutch nuclear families are generated by in vitro fertilization of fish collected from River Tyne. Six ♀MAR x ♂MAR families (red), six ♀FRESH x ♂FRESH families (blue), three F1 hybrid families from ♀MAR x ♂FRESH F1 hybrids and three reciprocal F1 hybrid families (purple). In each family two parents and nearly 94 offspring are whole-genome sequenced to more than 38X coverage for parents and >10X coverage for each offspring. b) For each nuclear family, paternal and maternal chromosomes of all offspring are phased separately. Switches from first haplotype to second haplotype (dark to light orange in maternal chromosome and dark to light green in paternal chromosome) are recorded as crossover (CO) events; defined as interval between last SNP of the first haplotype and first SNP of the second haplotype. c) Distribution of such crossover intervals across the chromosome is used to build a paternal and a maternal recombination map. d) Genome wide crossover count per meiotic product identified from individuals of each ecotype and sex shows consistently higher crossover count in females than in males. Error bar represents standard deviation. Mean crossover count is significantly reduced in hybrid females (marked as b) compared to mean of marine and freshwater females together (marked as a) e) Distribution of female (orange) and male (green) crossover events, plotted in 100 kb sliding windows across all 21 chromosomes, shows a more pronounced periphery-biased distribution in males than in females. Crossovers overlapping scaffold gap boundaries are excluded. Approximate position of centromere in all but three chromosomes (chrII, chrIV, and chrVIII) are plotted as vertical black dotted lines.

Our genetic map shows a high sex dimorphism, with a map length of 1081 cM for males and 1906 cM for females. Female sticklebacks produce significantly more crossovers per meiotic product than males (Fig 1d; mean in females: 19.06 ± 0.34 vs. males: 10.81 ± 0.09; *P* < 8 x10^-16^), an effect seen also in previous stickleback studies (9, 49). Half of paternal chromosomes (50.0%) and 68.6% of maternal chromosomes passed on to offspring have at least one crossover. While female-biased heterochiasmy is found across numerous taxa, e.g., *Drosophila* where no recombination occurs in males, and in several fishes (7, 50), the strength of heterochiasmy in sticklebacks (1.76 times more crossover per meiosis in females than in males) makes them a outlier among fish species (based on data summarized in (50)).

The genomic distribution of crossovers shows a striking overall bias towards chromosome peripheries and this effect is particularly strong in males (Fig 1e). More than 70% of male crossovers occur within the first or last 15% of the chromosome. Female crossovers have a more moderate periphery-bias (47%). On the 18 chromosomes for which approximate centromere location is known (51), acrocentric chromosomes tend to concentrate male recombination at the end of the long chromosome arm, except for chrXIX (the sex chromosome) for which males are the heterogametic sex and recombination between X and Y occurs in the pseudo-autosomal region (PAR).

We used forward simulations of three linked loci in populations undergoing adaptive divergence with gene flow to investigate the possible effects of heterochiasmy on the spread and selective sweep of adaptive mutations. Comparing strong heterochiasmy to a scenario with no heterochiasmy while keeping the overall recombination rate constant, we compared how quickly freshwater adaptive alleles spread through one marine-freshwater hybrid zone to a second freshwater population (detailed protocol in Supplementary methods 1). We found that increased variance in recombination rate as a result of heterochiasmy had little effect on the spread of adaptive alleles (*cf.* Fig S6 with Fig S8), but the magnitude of divergence between marine and freshwater populations was significantly higher in the presence of heterochiasmy (generation 120 Locus1 Δp_Pop1-Pop3_ = 0.79 +/-0.04 SD with hetererochiasmy, simulation 1, compared to Locus 1 Δp_Pop1-Pop3_ = 0.66 +/-0.06 SD without hetererochiasmy, simulation 3). We next contrasted a scenario where both males and females had high recombination rates to one where recombination was suppressed in males, and found recombination suppression in males marginally increased the probability and speed of a selective sweep and spread of adaptive alleles to new populations (*cf.* Simulation 3 to Simulation 5, Fig S10 and Fig S8 respectively).

### Genome-wide recombination reduction in hybrid females

While recombination variation between sexes is a widely observed phenomenon (4, 7, 9, 52, 53), some studies have also reported variation between closely related species or diverging ecotypes (11, 45, 54). At a genome-wide level, we find that marine and freshwater stickleback ecotypes do not differ in overall number of crossovers quantified directly from progeny, but a significant reduction in overall recombination rate is seen in hybrid females, compared to females of marine and freshwater ecotype (mean number of crossovers in hybrid females: 18.169±0.4SE; mean number of crossovers in marine and freshwater females: 19.5±0.42; Wilcoxon rank sum test, W=58, p=0.042, Fig 1d). This effect is also illustrated by total genetic map length (mean map length of 1816 cM in hybrid females versus 1950 cM in pure forms).

### Ecotype and sex effects on recombination rate

Since the local genomic actions of a recombination modifier may not be detected by genome-wide analyses, we used linear modeling to identify windows (500 kb) in which sex and/or ecotype can explain significant variation in crossovers. Ecotype explained significant amounts of variation in recombination rate in 6.75% of the genome (∼31.25Mb; Fig 2a), independent of sex, and with no clear bias in the direction or size of the effect. Marine and freshwater fish tended to have higher recombination rate in roughly the same number of windows (black points in Fig 2c). In contrast to ecotype, sex explained significant amounts of variation in recombination rate in 53% of the genome (∼245Mb; Fig 2a). This effect was independent of ecotype and, as expected, showed a clear bias towards higher recombination rate in females, with effects size of up to 10.23cM, while male-biased windows showed effects sizes of up to 8.87cM in PAR region and 5.9cM in autosomes (black points, Fig 2b). Interestingly, recombination variation among individuals in 15% of windows could be explained by an ecotype difference that was dependent on sex (sex*ecotype). Here, when marine and freshwater males differed in recombination, we observed that freshwater males had a higher recombination rate than marine males in most windows (green points; Fig 2c) – an effect that cannot be attributed to differences in detection power since nucleotide variation in these regions does not differ significantly between marine and freshwater males. In contrast, when ecotype explained significant variation among marine and freshwater females, the effect was reversed – marine females tended to have higher recombination rate than freshwater females (orange points; Fig 2c). It is notable though, that the maximum ecotype effect size was observed in a genomic window with freshwater female recombination rate being 5.57cM higher than that of marine females.

**Fig 2:**
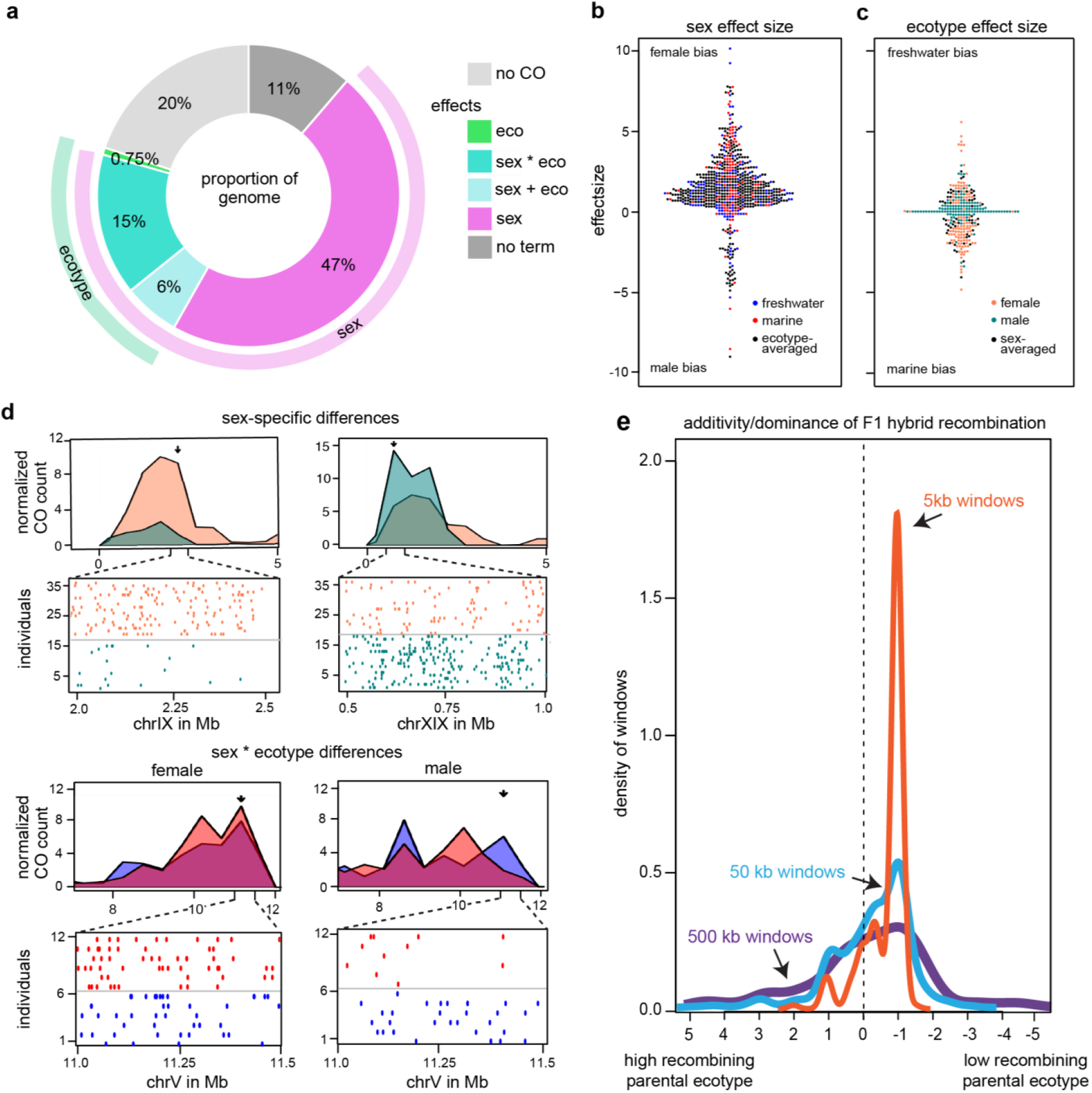
Sex of the individual explains three times more inter-individual variation in local genomic recombination than ecotype. a) Percentage of genome in which the individual’s recombination rate is influenced by sex and/or ecotype, determined by linear modeling on recombination rate in 500kb sliding windows (CO=crossover, eco=ecotype). b) Effect size (magnitude of difference in recombination rate) in local genomic windows where sex explains significant differences in recombination crossover count (black: sex-effect observed in both ecotypes, red: sex-effect only observed in marine ecotypes, blue: sex-effect only observed in freshwater ecotypes). c) Effect size (magnitude of difference in recombination rate) in local genomic windows where ecotype explains significant differences in recombination crossover count (black: ecotype-effect observed in both sexes, green: ecotype-effect observed only in males, orange: ecotype effect observed only in females). d) Examples of windows with significant differences in crossover counts (maximum effect size in plotted windows denoted with arrow). In the top panel, sex-specific crossover counts (orange: female, green: male) in a 5Mb region of chrIX (top left) and chrXIX (top right) are shown, with a zoom in of crossover midpoints below. In the lower panel, ecotype specific crossover counts (blue: freshwater, red: marine) in a 5Mb region of chrV, with a zoom in of a window exhibiting sexually antagonistic bias (marine-biased in females: bottom left, freshwater-biased in males: bottom right). e) Density estimates of dominance-additivity ratio of recombination crossovers in F1 hybrid relative to pure ecotype parental crossovers in genomic windows showing ecotype divergent recombination. A value of zero indicates recombination behaves additively in F1s relative to parents, 1 dominantly, -1 recessively, and values >1 or <-1 imply over or underdominance of recombination respectively. F1 recombination rate is biased towards the low recombining parental ecotype at all three (500kb, 50kb and 5kb) scales of resolution.

### Partially recessive recombination rates in hybrids

Using families with F1 hybrid parents, and considering only windows with significant differences among ecotypes, we asked whether recombination rate behaved in a dominant or additive manner. We found F1 hybrid recombination rate to be biased towards that of the low recombining parental ecotype irrespective of whether that parent was marine or freshwater. We hypothesize that modifier mechanism(s) suppressing local genomic recombination rate in both ecotypes behave in a partially recessive manner. This effect was observed in a sex-independent manner at broad as well as fine scale resolution (Fig 2e, Fig S2).

### Fine-scale recombination hotspots and coldspots

The genomic recombination landscape in sticklebacks was significantly non-random, with 80% of the crossovers occurring in less than 35% of the genome, and with the genome containing both ‗hot-‗ and ‗coldspots‘. The heterogeneity in the genomic distribution of crossovers (Gini-coefficient) in sticklebacks is intermediate to species that constrain most of their recombination into tight hotspots via the PRDM9 pathway (e.g. mouse (4) and human (5, 25)), and species that lack the PRDM9 pathway and have either less intense hotspots (yeast (16)) or no fine-scale hotspots (Drosopophila (13) and C. elegans (14); Fig 3a).

**Fig 3:**
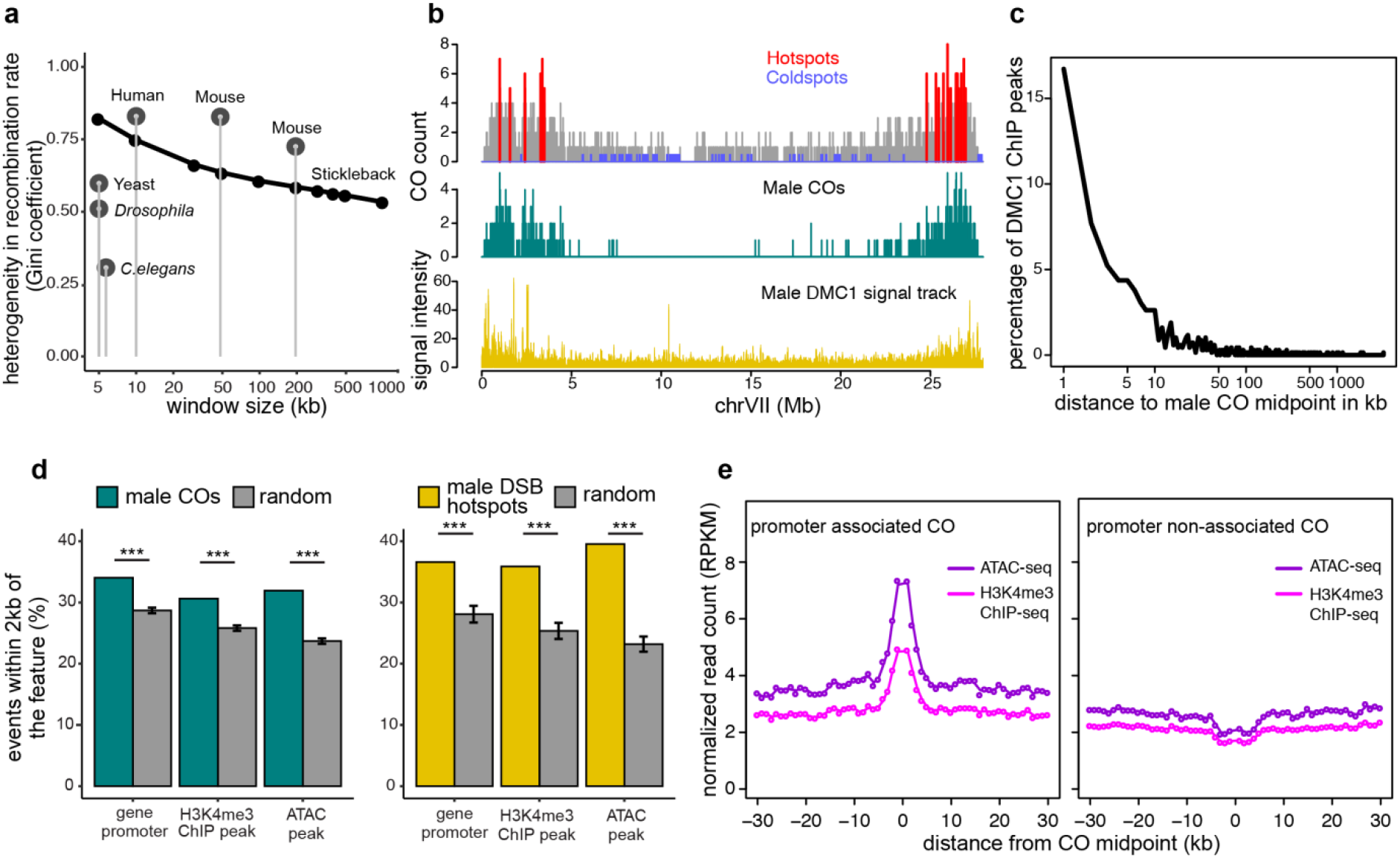
Molecular mechanisms and genomic features shaping the stickleback recombination landscape. a) Comparison of the heterogeneity (Gini-coefficient) of the crossover distribution in the genomes of human (5), mouse (4), yeast (16), Drosophila (56), and C. elegans (14) at the scales the studies were done in (grey dots), and the stickleback data of this study at various scales (black line). b) The crossover landscape (grey peaks) contains recombination hotspots (red peaks) and coldspots (blue peaks). The overall shape of the sex-averaged recombination landscape is highly similar to the DSB landscape, as mapped by anti-DMC1-ChIP-seq in males (bottom panel, yellow peaks). c) In stickleback males, distances from anti-DMC1-ChIP-seq signal peaks (proxy for DSB hotspots) to the nearest crossover shows that not all crossovers take place in the proximity of a DSB hotspot. d) In stickleback males, crossovers (left panel) and DSB hotspots (right panel) are significantly associated with gene promoters, with H3K4me3 histone modification marks, and with chromatin accessibility (assayed with ATAC-seq). e) In stickleback males, promoter associated crossovers coincide with elevated signals of chromatin accessibility (purple) and H3K4me3 (pink) (left panel). On the contrary, crossovers further than 2 kb away from the nearest promoter coincide with decreased signals of chromatin accessibility and H3K4me3 (right panel).

We defined stickleback hotspots as intervals containing multiple crossovers (at 5% false discovery rate (FDR); Fig 3b). At a fine-scale of 5kb intervals, the crossover rate within hotspots was on average 3.7 times more than that of the flanking region and ten times more than the genome-wide average, with an intensity ranging from 0.15 to 0.45 cM. Stickleback hotspots are six times less intense than mouse and human (0.001cM to 3cM; (55)) and nearly five times less abundant (after accounting for differences in genome size (4)). Pooling crossover counts within ecotypes and considering the genomic colocalization of observed crossovers from marine-, freshwater- and hybrid-ecotype fish we identified 12 distinct genomic regions (comprised of 31 overlapping 5kb windows) with significant log_2_ fold change differences in crossover counts in pairwise comparisons among ecotypes (5% FDR; Tables S2-S5). We observed more freshwater ‗hotspot‘ regions than marine and hybrid (*cf.* sixteen, eight, and seven 5kb windows respectively, and seven, three, and three genomic regions respectively), with the two strongest ecotype-divergent 5kb hotspot windows containing log_2_ fold differences in crossover count between freshwater and hybrid individuals >=6.34 (chrXXI:11400304–11405304), and >=6.34 between marine and hybrids (within the pseudoautosomal region of the X chromosome chrXIX:1625740–1630740). All 31 of the hotspot windows identified contained crossovers from 5–12 fish, suggesting the signature of ‗hotness‘ was not driven by recombination patterns within a single individual. The majority of hotspots were exclusively ‗hot‘ in only one ecotype, and at hotspots differing between marine and freshwater ecotypes, F1 hybrids showed crossover counts that were equal to or marginally higher than the ‗cold‘ ecotype (Tables S3-S5).

Defining recombination coldspots as contiguous regions that could accommodate at least five crossovers assuming a uniform distribution, but contained zero crossovers, we identified 499 distinct coldspots with a minimum size of 212kb. Whereas hotspots were most often located in sub-telomeric regions of chromosomes, coldspots were most often located in the centers or telomeric ends of chromosomes (Fig 3b). In females, we found hybrid coldspots to be significantly larger (median 345kb) than freshwater (323kb) or marine coldspots (311kb) (Wilcoxon rank sum tests, W=20423 and 24869; p=0.027 and 0.023 respectively). This effect was not significant in males.

### Recombination is shaped by location of DNA double-strand breaks and gene features

Regulation of crossover location in meiosis may occur via ‗upstream‘ mechanisms determining the location of DNA double strand breaks (DSBs) or alternatively via ‗downstream‘ mechanisms influencing whether DSBs are repaired as crossovers or non-crossovers. On the molecular level, chromatin accessibility and histone modifications such as H3K4me3 are known to attract DSBs (reviewed in (57)), that can be repaired either as allele-shuffling crossovers or as non-reciprocal non-crossovers ((58, 59)). To determine the importance of ‗upstream‘ versus ‗downstream‘ mechanisms in shaping the genomic recombination landscape in sticklebacks, we generated a DSB map in male meiotic cells and compared this to male crossover maps generated from nuclear families (above). We produced an antibody against stickleback meiotic protein DMC1, that binds to the single-strand ends of meiotic DSBs, and performed chromatin immunoprecipitation followed by sequencing using a modified protocol for recovery of single-stranded DNA-protein interactions (60) (anti-stickleback DMC1 ChIP-seq) on testes tissue, using liver tissue as a negative control. Testes-specific DMC1-signal was elevated at the chromosome peripheries, in a pattern highly similar to the genomic distribution of male crossovers (Fig 3b). We identified a total of 1090 signal peaks with median size of 1.4kb (indicative of DSB hotspots), 72.2% of which were located in the first or last 15% of the individual chromosomes, consistent with the distribution of crossovers inferred from nuclear families above (70%). Further, we see prominent proximity enrichment of testes-specific DMC1-signal peaks to male crossovers identified from nuclear families (38.7% of DMC1 peaks have a male crossover within 5kb distance, despite different individuals being used in the distinct assays; Fig 3c). This enables a lower estimate that two-fifths of stickleback male DSB hotspots are associated with repair via crossovers while the remaining three fifths of DSBs may repair via the non-crossover resolution pathway. We anticipate that the estimated proportion of DSBs resolved as crossovers would increase if it were technically possible to use meiotic cells from the same male individual in both assays. Combined, these results suggest that in sticklebacks the genomic recombination landscape is shaped by mechanisms determining the location of DSB initiation.

In other species, recombination has been found to often coincide with genomic features such as promoters and repetitive regions (19, 60), and in many mammals with specific DNA motifs recognised by trans-acting mediators of recombination, e.g., methyltransferase PRDM9. To further understand the molecular and genomic factors regulating crossover distribution in sticklebacks, we tested the association of crossovers with chromatin accessibility in meiotic cells (stickleback primary spermatocytes); gene features; active promoters predicted by the histone modification H3K4me3 from ChIP-seq on stickleback testes; and DNA motifs. Male crossovers identified from nuclear families, but not female crossovers, were significantly associated with genic regions and all genic features (most strongly with promoters, but also with introns and exons, and more weakly with transcription end sites). Sticklebacks lack a homolog to the mammalian hotspot mediator *PRDM9*, and we found no evidence of motifs enriched in crossover hotspots aside from short CpG-like GC rich motifs significantly correlated with crossover hotness (Table S6). Predicted DSB hotspots (DMC1-ChIPseq signal peaks) from male testes tissue, as well as male crossovers from nuclear families, were proximally associated with regions of active transcription (Fig 3d; predicted by chromatin accessibility at promoters, and the histone modification mark H3K4me3). 46.8% of crossovers and 54.2% of predicted DSB hotspots were found within 2kb of active transcription while the remaining half of male recombination crossovers were more distal to these features. Since the crossover and DSB datasets are derived from different individuals it is possible these more distal crossovers are associated with active transcription not detected in individuals used in DSB assays. However, we note with interest that crossovers occurring further than 2kb away from the nearest promoter colocalize with a decrease in chromatin accessibility and H3K4me3 marks (Fig. 3e) suggesting they are enriched in regions of inactive transcription. These signatures might be indicative of an undescribed means of recombination regulation away from gene features in stickleback fish.

### Adaptive ‘islands’ fall in regions of low recombination

Next, we considered the evolutionary implications of differences in recombination rate in the context of adaptive loci. When adaptive divergence evolves in the face of gene flow, the shuffling of adaptive loci can lead to mismatched combinations of divergently adaptive alleles, which may have important fitness effects on the individual and ultimately populations. Adaptive divergence of freshwater stickleback ecotypes from marine ecotypes draws strongly on standing variation in the form of highly divergent marine–freshwater haplotypes (up to 4% sequence divergence) at 242 genomic loci (FDR 0.05), including three large inversions (32). On a broad genomic scale, loci underlying adaptive divergence of marine and freshwater fish fall in regions of low recombination rate (e.g., Fig 4a; see also (34, 42, 43)). Defining adaptive islands as chromosomal regions of linked adaptive alleles, we found recombination rate in males to be nearly three times lower inside adaptive islands compared to outside (Fig 4 a,b). Females show a similar pattern though the magnitude of effect is smaller (1.2 times lower inside than outside; Fig 4 a,b). Using whole-genome sequencing data to determine the proportion of marine genome composition in each of the individuals studied (see methods), we asked whether the recombination rate within and outside ‗adaptive islands‘ differs among individuals depending on their genome-wide admixture proportion using a non-linear quadratic regression. We observed a tendency for individuals with admixed genome composition to have a significantly lower recombination rate outside adaptive islands, compared individuals with either freshwater or marine ancestry (Fig 4b).

**Fig 4:**
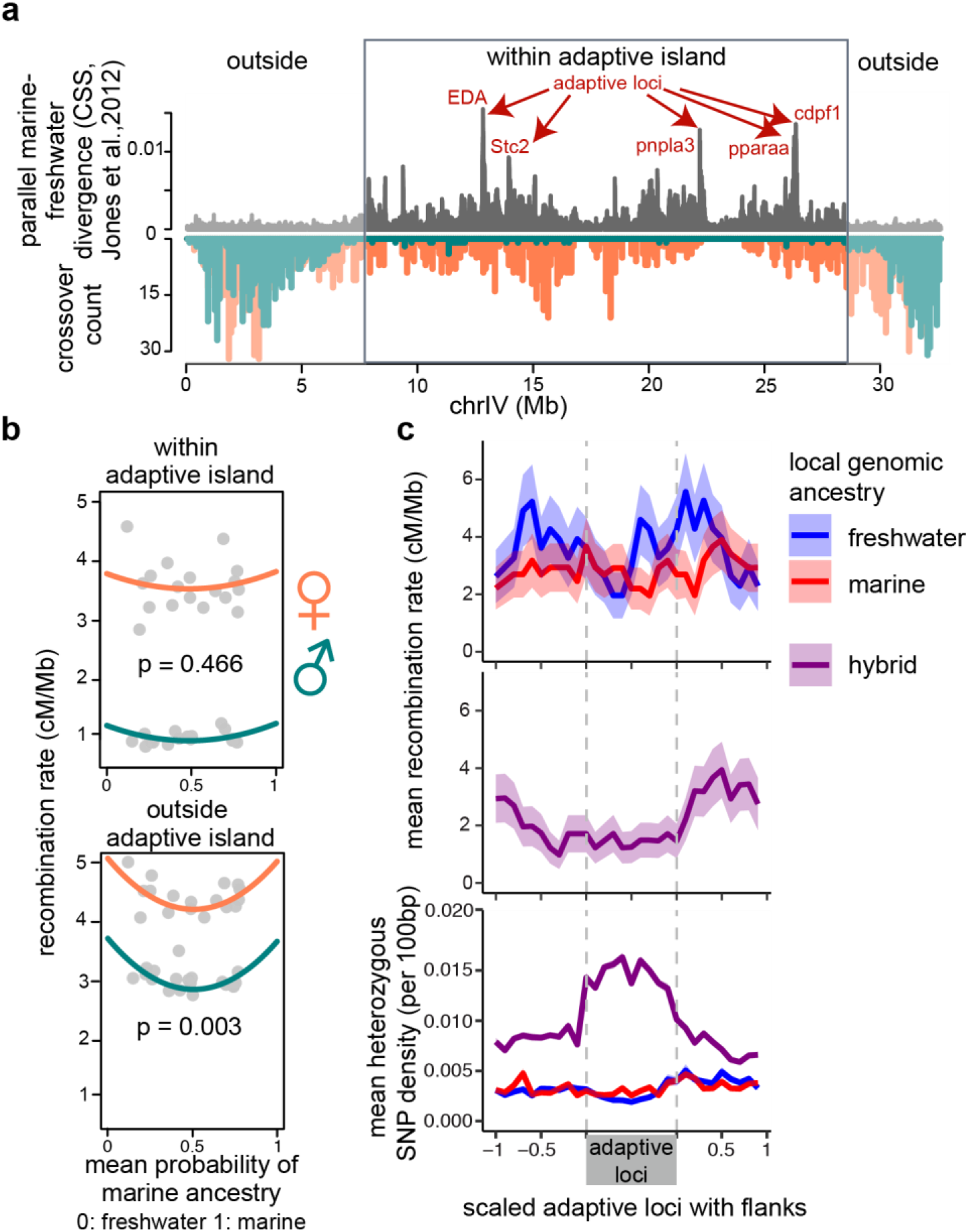
Recombination suppression within and among adaptive loci. a) Parallel adaptive divergence (grey, cluster separation score from Jones et al. 2012 (32)) and sex-specific crossover counts in 100kb sliding windows (male: green, female: orange) show a suppression of male crossovers, and to a lesser extent of female crossovers, within an adaptive island on ChrIV. b) Recombination rate within adaptive islands is lower than outside adaptive islands in both sexes. The strength of recombination suppression outside, but not inside adaptive islands seem to correlate with the degree of genetic admixture. c) Admixed genetic ancestry at focal adaptive loci (heterozygous for marine and freshwater haplotypes, purple) is associated with lower recombination rate within the divergent adaptive haplotypes, and elevated recombination rate on the divergent haplotype flanks (top panel). Mean female recombination rate (and standard error) across these 242 adaptive regions grouped by their local ancestry (top panel - homozygous marine: red and homozygous freshwater: blue; middle panel - heterozygous: purple) shows reduced recombination in loci with hybrid ancestry. Mean heterozygous SNP density in 100bp windows across the scaled adaptive loci in the same categories shows a high heterozygous SNP density in adaptive loci with hybrid ancestry (bottom panel).

### Recombination suppressing effects of heterozygosity

The occurrence of female crossovers inside adaptive islands allowed us to test whether recombination crossovers shuffle, disrupt and shape the boundaries of divergent haplotype blocks at individual adaptive loci. In contrast to individuals with marine or freshwater genomic composition, individuals who were heterozygous for marine and freshwater haplotypes at adaptive loci show changes in recombination rate that follow the boundaries of the divergent adaptive loci – lower recombination within adaptive loci, and elevated recombination on the flanks (Fig 4c). Associated with this boundary effect in these individuals, heterozygosity (mismatches between chromosome homologs) is elevated within adaptive loci (Figure 4c) and lower in the flanks. Here, we avoid any bias from differences in population history, because the observed suppression in recombination is based on crossovers identified from pedigrees and so is not confounded by changes in LD decay around loci subject to selection, nor by contrasting demographic histories of the different ecotypes that confound comparisons of *rho*. The association between elevated heterozygosity and low recombination is consistent with the observations of recombination suppression in inversion heterokaryotypes (Fig S3; (61)), and colocalization of recombination coldspots with high heterozygosity in genomic regions of otherwise high recombination rate (e.g. Fig S4a). Across the entire dataset we note that the vast majority of crossovers occur in regions of low heterozygosity (less than 50 heterozygous sites per 5kb, approximately one heterozygous site per 100bp; Fig S4b). We hypothesise that high heterozygosity slows the rate of homolog engagement during double strand break repair and increases the likelihood of repair by non-crossover pathway (62, 63). This phenomenon may have important implications for populations undergoing adaptive divergence with gene flow – using forward simulations of three linked loci in populations undergoing adaptive divergence with gene flow, we found that recombination suppression in cis(coupled)-heterozygotes increases the rate at which freshwater adaptive alleles spread through marine-freshwater hybrid zones and are swept to fixation in new freshwater populations (Fig S6-S13). Further, under a selection migration balance, recombination suppression in cis(coupled)-heterozygotes helps to maintain significantly higher divergence in allele frequencies between marine and freshwater populations (for example, compare Simulations 1 and 2 Fig S6 and Fig7 respectively, Table S8).

### Fitness effects of recombination in natural populations

Quantifying the fitness effects of recombination in wild populations is challenging, because it requires tracking fitness-affecting loci (and recombinants) in a large number of individuals. Stickleback chromosome IV harbors a number of loci underlying variation in fitness (64), major skeletal traits (36, 65) and other pleiotropic effects (66), and includes five genomic regions with extraordinarily strong molecular signals of parallel marine-freshwater divergence (Fig 3a; *EDA* chrIV:12.81, *Stc2* chrIV:13.94, *pnpla3* chrIV:22.21, *pparaa* chrIV:26.30, *cdpf1* chrIV:26.35,; (32)). Although these five loci span a large physical distance (more than 13.5Mb), the low recombination rate (1.31cM/Mb) of this region maintains strong linkage among adaptive alleles and facilitates their inheritance as a linked adaptive cassette. Furthermore, the strength of selection acting on this region in newly founded freshwater populations is extremely strong (contemporary evolution estimates of selection coefficient at EDA and its linked genomic loci *s*\*=0.5; (64)). We hypothesized that when gene flow occurs between divergently adapted populations, the shuffling effects of recombination among adaptive loci may have deleterious consequences for the fitness and survival of admixed offspring.

We leveraged a previously described natural marine-freshwater hybrid zone with large population sizes to explore the potential fitness effects of recombination in the wild (67) in two ways: Firstly, using standard length as a proxy for fitness (ability to grow), and secondly using changes in within-chromosome IV two-locus LD over temporal samples of the same cohort as an indication of differential survival of recombinant genotypes. 285 young-of-the-year from the hybrid zone in River Tyne, Scotland (Fig 5a) (67) were finclipped, tagged, and released and their DNA genotyped using a 3084 SNP genome-wide custom genotyping array. Focusing on chromosome IV, we calculated the minimum number of parental recombination events needed to explain an individual‘s multilocus genotype at the five linked loci (Fig 5b,c). This is distinct from a ‗hybrid index‘ that is commonly used in hybrid zone studies, since potential F1 hybrids heterozygous at all five loci are assumed to be carrying non-recombined marine and freshwater chromosomes and are assigned a score of zero, while the maximum possible number of recombination events between the five loci on the two chromosome copies is eight and would be achieved when an individual alternates between homozygous marine and freshwater genotypes across the five loci.

**Fig 5.**
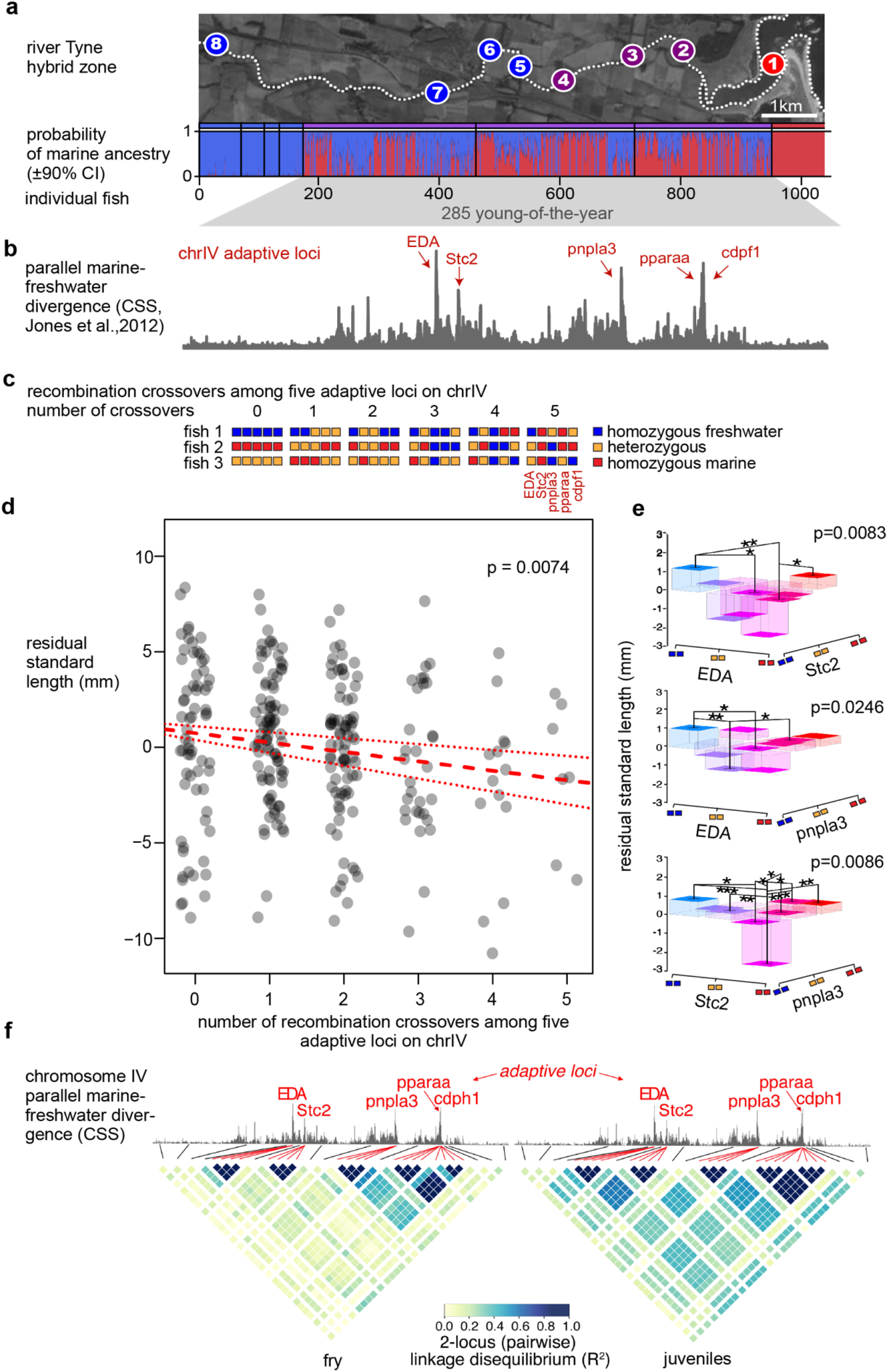
Selection against hybrid recombinants in a natural hybrid zone. a) Sampling sites at the hybrid zone in River Tyne, Scotland, and probability of marine ancestry of sampled individuals (red: marine, blue: freshwater). b) Parallel genetic divergence between global marine and global freshwater populations (CSS, from Jones et al 2012 (32)). c) Minimal cross over counts among five adaptive loci on chromosome IV were calculated from multilocus genotypes with example genotypes for each count shown. d) Body size (residual standard length, a proxy for fitness that reflects size relative to cohort mean), decreases as the number of recombination events between adaptive loci on chromosome IV increases. d and e) Individuals carrying mismatched marine and freshwater alleles across Eda:Stc2, EDA:pnpla3, Stc2:pnpla3, Stc2:cdpf1, Stc2:pparaa loci are significantly smaller in size than individuals with non-mismatched combinations of alleles. This effect is significant both as a (d) simple linear count of alleles and (e) under an epistasis model between pairs of loci. f) Linkage disequilibrium among adaptive loci increases as a cohort ages – a pattern consistent with selection against recombinants.

We observed a significant negative relationship between body size and the minimum number of recombination events an individual carries among adaptive loci on their diploid chromosomes IV (effect size slope -0.492+/-0.182SE mm/recombination event, p=0.0074; Fig 5d). Recombinant individuals carrying a mixture of marine, freshwater and/or heterozygote genotypes at the five adaptive loci were smaller than individuals with fewer recombination events, suggesting these individuals grow less well than non-recombinants – an effect that cannot be explained by single locus genotypes at the five individual adaptive loci alone (all p-values >0.05). Rather, consistent with the importance of a linked adaptive cassette, we find evidence for epistasis among adaptive loci on this chromosome; linear modeling revealed that two-locus genotypes at EDA:Stc2, EDA:pnpla3, Stc2:pnpla3, Stc2:cdpf1, Stc2:pparaa explained significant amounts of variation in standard length (anova F_1,281_ range 4.56-7.07, p=0.0083-0.0337, Fig 5e). Individuals with mismatched two-locus genotypes (e.g. MM:FF; MM:FM, FM:FF) were significantly smaller in body size than MM:MM and FF:FF genotypes (posthoc ttest p-values range <0.001-<0.05)(Fig 5e). Carrying recombinant copies of chromosome IV may also reduce the probability of survival: When examining changes in genotype composition of the same cohort over time, we observed significantly higher minimum crossover counts among the five adaptive loci in fry (sampled in July & August) compared to juveniles sampled 2-3 months later (September & October; mean crossover number in fry=1.845, mean crossover number in juveniles=1.313; t_233.93_=3.675, *P* < 0.0003). Consistent with this we found that pairwise linkage disequilibrium among adaptive loci strengthened as the cohort aged from fry to juveniles (Fig 5f). In the older juveniles cohort, an individual‘s genotype at an adaptive locus was more predictive of the alleles it carried at other adaptive loci, an effect that may be caused by higher mortality of recombinant individuals over time. Combined, these results suggest that individuals carrying mismatched marine and freshwater alleles caused by recombination among adaptive loci on chromosome IV grow and survive less well than their non-recombinant peers.

## Discussion

Linking recombination variation to its evolutionary consequences in natural populations is a major challenge. By constructing individualized high-resolution one-generation crossover maps in an evolutionary model organism, the threespine stickleback fish (Fig 1), our empirical recombination study enables us to establish ecotype- and sex-specific estimates of recombination rate unconfounded by the varying effects of demography and selection in different populations. This perspective complements previous approaches that have used sex-averaged population genetic estimates such as rho (15, 45, 68–70). To further investigate the molecular basis of recombination regulation, we juxtapose our sequencing-based maps with findings of the molecular signatures at crossovers as well as double strand break (DSB) hotspots. Finally, using a field study of a natural marine-freshwater hybrid zone, we provide empirical evidence indicating deleterious fitness consequences of recombination.

Sex-differences in recombination are commonly observed across diverse organisms. Among fishes, species with male- and female-biases are equally common (50). Our study of crossovers inferred from large nuclear families show a strong female-bias with 1.76 times more recombination events in females than males (Fig 1d) and is consistent with previously reported studies in sticklebacks (9, 49). Approximately half of the chromosomes in a male stickleback gamete are inherited faithfully without a crossover. This can be explained by a single crossover between two of the four sister chromatids making up a bivalent homologous chromosome pair, followed by independent segregation of these chromatids (2 with crossovers, 2 without) into four haploid gametes. Since at least one crossover is necessary to ensure stable pairing and proper segregation of chromosome homologs it is likely this represents a mechanistic constraint that imposes a lower limit on recombination rate. Given stickleback males recombine closer to this lower limit, evolution of further recombination suppression is likely constrained. In contrast females have considerably more crossovers, and are therefore less constrained by lower mechanistic limits and are able to lower recombination rates without segregation failure. This provides a plausible explanation as to why we detected significant recombination suppression in female hybrids but not male hybrids.

We find a location bias towards the sub-telomeric regions of the genome (Fig 1e), and that these regions show signatures of long-standing elevated recombination rate, such as increases in nucleotide diversity and GC content (Fig S5). While haploid selection against male gametes with crossovers in the middle of chromosomes cannot be fully excluded, our previous studies of empirical recombination crossovers in male sperm show a similar sub-telomeric bias (61). This sub-telomere bias is generally attributed to conserved features and processes of meiosis, such as chromosome condensation, centromere position, telomere bouquet formation and crossover interference (19, 60, 71, 72). Interestingly, gene density is uniform across the length of the chromosomes (Fig S5) but loci underlying parallel divergent adaptation (32) are found only in the recombination-reduced inner regions suggesting that the broad scale recombination landscape has shaped the genetic basis of ecotype divergence.

We applied linear modeling to quantify the effects of sex and ecotype on recombination rate, and show that although sex is the major predictor, also ecotype and the interaction of sex and ecotype affect more than 20% of the genome (Fig 2 a,b). The direction of ecotype effect was balanced between marine and freshwater, but did have a strong female-bias (Fig 2b), as expected from the overall higher frequency of crossovers in females. Comparing the local recombination rates between parental ecotypes and hybrid offspring revealed the ecotype effect to be partially recessive (Fig 2e). Importantly, we found that this tendency to recapitulate the recombination rate of the lower recombining parent was not biased towards either of the parental ecotypes or sexes. This supports a role for divergence in cis-acting recombination modifiers whose effects are disrupted in heterozygotes/hybrids.

Our results raised the question how precisely recombination location is controlled in sticklebacks. Organisms lacking the chromatin modifier PRDM9 are thought to represent the ancestral state of recombination regulation, in which double-strand breaks (DSB) occur preferentially in accessible chromatin, leading to a conserved pattern in crossover distribution (44, 73, 74). Despite sticklebacks lacking PRDM9, their crossover distribution is still non-random and draws nearer to species with intense PRDM9-mediated hotspots (Fig 3a). Using ChIP-seq against DMC1 in male testes containing meiotic cells, we were able to confirm that these ‗semi-hot‘ hotspots do recapitulate the pattern of DSB hotspots (Fig 3b, c). This provides strong evidence for stickleback crossover frequency being regulated by factors influencing DSB initiation. Additionally, we found two classes of crossovers: promoter-associated crossovers that are strongly coinciding with open chromatin and the activating histone mark H3K4me3, and promoter-distal crossovers occurring far away from promoters are associated with a small but detectable reduction in chromatin accessibility (Fig 3e). Further investigation is needed to determine the mechanisms influencing DSB formation in this latter class of crossovers, though active nucleosome remodeling, involving proteins other than PRDM9 may be required.

Taking together our findings of partial recessivity and crossover hotspot regulation, we asked if recombination is suppressed in ecotype hybrids within the divergently adaptive loci. We found indeed that adaptive alleles are shuffled at an exceptionally low rate in hybrid females, and hardly at all in hybrid males. We also observed that the recombination rate is lower in local genomic regions of high heterozygosity (Fig 4a, b, c). Variance in recombination introduced via heterochiasmy and recombination suppression in heterozygotes may have evolutionary implications for populations undergoing adaptive divergence in the face of gene flow. Our forward simulations revealed that when new adaptive alleles arrive in linkage with other beneficial mutations (a plausible scenario when the genomic basis of parallel adaptative divergence is highly polygenic), heterochiasmy can help maintain greater divergence between populations (Fig S6 and S8, Table S8), while recombination suppression in cis(coupled)-heterozygotes can increase the rate at which new beneficial mutations establish and spread through populations maintained in selection-migration balance.

We explored the potential fitness implications of recombination in the wild, by following the growth and genetic makeup of a cohort of young-of-the year across a marine-freshwater hybrid zone (Fig 5a). We found allelic shuffling among five loci underlying adaptive divergence to come with a cost to growth and survival (Fig 5b-f) and hypothesize that this deleterious fitness effect may form the basis by which natural selection has favored recombination modifiers that suppress recombination in hybrids and heterozygotes. Combined, our results highlight how both molecular constraints and population-level evolutionary forces have the potential to shape the recombination landscape.

Leveraging multiple study approaches on stickleback fish populations undergoing adaptive divergence in the face of gene flow, we have quantified fine-scale variation in recombination across the genome, shown evidence and possible molecular mechanisms for recombination suppression in hybrids, and the fitness consequences of recombination among divergent ecotypes in the wild. Our results support predictions that divergence in cis-acting recombination modifiers whose mechanisms are disrupted at local genomic regions of heterozygosity and in hybrids across the entire genome may have an important role to play in the maintenance of differences among adaptively diverging populations. Further studies will be needed to address the role of the strong sex-bias in recombination rate, and to illustrate the molecular mechanisms through which natural selection shapes recombination.

## Materials and methods

### Ethics statement

ll animal experiments were done in accordance with regulations of the U and of the state of aden-W rttemberg, ermany. The ax lanck Society holds the permits to capture and raise sticklebacks. The stickleback fish facility is maintained under aden-W rttemberg regional authority permission (ompetent authority: egierungspraesidium T bingen, ermany; ermit and notice numbers 35 9185.82-5, 35/9185.46)

### Generating data for individualized crossover maps

Stickleback fish were collected from River Tyne in Scotland. Marine and freshwater sticklebacks were caught from 4 km and 19 km respectively from the river mouth during May-June 2014. Following the standard protocol, in vitro fertilization crosses were carried out in the field and the embryos were raised in a stickleback fish facility at the ax lanck campus in T bingen. oth marine and freshwater fish were raised in 10% seawater salinity (3.5 ppt) with daily 10% water change. All fishes were fed once a day with the same food that consists of both marine and freshwater invertebrate diets. For fine-scale individualized recombination map construction,18 nuclear families consisting of 90-94 offspring per family were generated by in vitro fertilization of 36 distinct parent individuals (6 marine ♀ X marine ♂ crosses, 6 freshwater ♀ X fresh water ♂ crosses, 3 (freshwater X marine) ♀ X (freshwater X marine) ♂ crosses, and 3 (marine X fresh water) ♀ X (marine X fresh water) ♂ crosses). Out of these 18 crosses, two freshwater and two marine crosses were carried out in the field (Family X1, Family X4, Family X11, Family X20) whereas 14 crosses were made in the lab using offspring grown in aquariums. For crosses that resulted in small clutch size, a second round of in vitro fertilization was carried out in the next breeding season with eggs from the same female and cryo-preserved sperm from the same male fish. DNA was extracted either from a piece of tail fin (parents) or from the whole fish body (offspring at age of one month) following Solid Phase Reverse Immobilization (SPRI) bead based Protocol (75). Multiplexed DNA extraction was carried out using a TECAN liquid handling robot. A home-made low-cost library preparation protocol, adapted for high throughput from (76) was followed for library preparation (detailed protocol in Supplementary methods 2). Libraries (with ∼250 bp insert size) were sequenced as 150bp paired-end reads on an Illumina HiSeq3000 sequencer to about 10X genome coverage per offspring in each family and about 60X coverage per parent (Table S1). Sequenced reads were mapped to the stickleback reference genome gasAcu1 (Broad S1 assembly generated from an Alaskan freshwater female) (32) using Burrows-Wheeler Aligner (BWA) v0.7.10-r789 (77) with bwa mem option. Mapped reads were then sorted and indexed using SAMtools (78). Further read processing until variant calling was carried out according to the best practices recommended by Genome Analysis Toolkit (GATK) (79, 80).

### Variant calling and filtering

Individual-wise variant calling was carried out using the HaplotypeCaller option in GATK v3.4. Individual gVCF files from a family were then combined using the GenotypeGVCF option in GATK v.3.7. This step creates a joint genotype file for a family with information of parents and all offspring at each variant position. High quality variants for further analysis were then selected based on the following criteria: 1) SNPs (indels are excluded), 2) biallelic, 3) heterozygous in either parent, 4) SNPs with quality score greater than first quantile of the quality score distribution of the whole family data set, 5) model-based clustering analysis of allele frequency was carried out using R package mclust version 5.4.1 (81). Since in each nuclear family, alleles segregate in mendelian ratio, SNPs falling in tight clusters of allele frequency around 0.25, 0.5 and 0.75 were selected, 6) SNP filtering based on GATK hard filtering criteria (QD < 2.0 || FS > 60.0 || MQ < 40.0 || MQRankSum < -12.5 || ReadPosRankSum < -8.0), 7) SNPs with coverage not differing by more than 50% of mean coverage and read bias between alleles less than 30% were selected (Table S1).

### Scaffold orientation correction

The stickleback reference genome, Broad S1 assembly (gasAcu1) was used for the initial mapping. However, orientation of 13 scaffolds in gasAcu1 has been corrected in an improved version of the assembly (47). Since the scaffold orientation can affect phasing and crossover identification, the order of SNPs in these 13 scaffolds was reversed prior to haplotype phasing using a custom made perl script. It has to be noted that only the orientation of scaffolds in the assembled part of the genome was corrected. In contrast to the latest version of the assembly, no new scaffolds were additionally tied into the assembled chromosomes from unassembled scaffolds, making our crossover counts a conserved underestimate.

### Haplotype Phasing

The phasing algorithm SHAPEIT (82) in combination with duoHMM (83)(O’Connell et al. 2014) was used to phase SNPs within a family. The joint vcf file for a family was split into a vcf file for paternally informative SNPs and another file for maternally informative SNPs. Paternal informative SNPs are those that are heterozygous in the father while being homozygous in the mother and vice versa for maternally informative SNPs (SNP counts in Table S1). SHAPEIT + duoHMM requires a genetic map for each chromosome as an input. In order to avoid any bias in phasing due to the preexisting recombination rate per window, a linear genetic map with 3 cM/Mb (reported genome-wide average recombination rate in sticklebacks is 3.11 cM/Mb (42)) recombination rate was used. The first round of phasing was then carried out with SHAPEIT without using any pedigree information. The SHAPEIT output files were then fed into duoHMM to identify and correct phasing errors by taking pedigree structure into consideration. In addition, duoHMM also generates a list of SNPs with genotyping error probability. After one round of SHAPEIT and duoHMM, problematic SNPs were short-listed based on the two following criteria;

1. SNPs with genotyping error probability >0.9 in 20 or more offspring
2. SNPs showing biased transmission distortion (phased to one haplotype in more than 75% of offspring). While some transmission bias may be biologically real due to processes like meiotic drive, it may also be bioinformatic error and therefore these SNPs were removed from the analysis. A second round of phasing (SHAPEIT + duoHMM) was then carried out by excluding those error-prone SNPs from the data set.

### Crossover identification

We developed an R based pipeline employing the following strategy to identify CO events: long switches (haplotype switches with >50 kb size on either side and 50 or more SNPs supporting each haplotype) in phased haplotypes were identified as true crossover events, filtering out all small switches that are either genotyping errors or gene conversion events. After calling crossover events from every sequenced offspring, further filters were applied on crossover list in order to remove false positive events detected as a result of phasing error or low sequencing coverage; 1) COs appearing in 50% or more offspring in a family between the same boundary SNPs (probably due to phasing error) were removed; 2) all COs from offspring who had sequencing coverage <2x were removed; 3) COs with low resolution (>1Mb) were removed unless due to the lack of informative SNPs; 4) COs at inversion boundaries were removed (further details of this filter can be seen in the section on inversions).

### Linear modelling of sex and ecotype variation

We performed linear modelling on every 500kb window across the genome considering pure form individual recombination rate as the response variable and sex, ecotype, or both in combination, as the explanatory variable using the linear regression function in R. Only windows with non-zero recombination rate in at least one individual were included in the analysis. Minimal model explaining the variation was obtained by applying the step function. 100,000 random shuffling of crossovers was performed to estimate false discovery rate of finding an effect in each window and windows with FDR<0.05 were shortlisted. In a window with significant influence of either sex and or ecotype, effect size was calculated by subtracting mean categorical recombination rate. For example, In a window with significant sex effect, mean male recombination rate was subtracted from mean female recombination rate to get the effect size and the direction of effect.

### Additivity/Dominance in F1 hybrids

In windows with ecotype effect (identified from linear modelling above) deviation in hybrid recombination rate from mid-parental recombination rate (mean recombination rate of parental ecotypes in the respective windows) was calculated (hybrid - midparental value). A value of zero indicates hybrid showing additivity whereas a positive value indicates dominance of high-recombining ecotype (recombination regulator acting dominantly) and negative value indicates dominance of low recombining ecotype (recombination regulator acting recessively) in hybrids. This analysis was performed at three different scales (5kb, 50kb, and 500kb) and in a sex independent and dependent manner.

### Genomic ancestry calculation from wgs data

We estimated the probability of marine, freshwater, or hybrid ancestry at each SNP position in each parental individual based on allele frequency estimated from 11 freshwater and 10 marine fish sequenced in Jones et al., 2012 (32) study. In the 21 genome data set, a genotype observed at high frequency in marine individuals was defined as marine genotype and the freshwater genotype was defined similarly. Probability of ancestry at each site in each parental genome of this study is assigned based on the number of reads covering the marine or freshwater allele. Mean ancestry probability in regions of interest such as adaptive loci and adaptive islands were then calculated and further correlated with recombination rate to investigate the effect of regional ancestral allele state on recombination.

### Recombination at adaptive loci

In order to allow comparison of recombination rate within and immediately flanking adaptive loci, we first rescaled 242 adaptive loci (32) to a fixed width and calculated mean recombination rate and standard deviation in sliding windows across the adaptive loci ± flanks. Adaptive loci were further grouped based on mean local genomic ancestry and repeated the analysis.

### Identifying candidate hotspots that differ among ecotypes

The 1bp midpoint of crossover intervals from each individual were pooled by ecotype and downsampled to create three equal sized sets of 9463 marine, freshwater and hybrid crossovers that excluded crossovers spanning scaffold boundaries and/or those with interval boundaries exceeding 10kb. The number of crossovers within 5kb of each midpoint was counted for all three ecotypes, and the log2 foldchange ratio of crossover counts was calculated for pairwise ecotype comparisons using a pseudocount of 0.1. Significance of the foldchange in crossover count being different from zero was determined by comparison to the upper and lower 0.025 tails of foldchange values from a null distribution, accounting for mean average crossover count (similar to MA analyses in analysis of differential gene expression). Specifically, this null distribution was created for each pairwise ecotype comparison by shuffling the downsampled ecotype crossover locations into two random sets of 9463 crossovers 10000 times, counting the number of crossovers in each set in 5kb windows and calculating the log2 foldchange ratio of crossover counts in pairwise comparisons among sets. 2.5% quantiles of the foldchange distribution for a given mean average number of crossovers were then calculated and used to identify observed foldchange ratios that would occur by chance with a probability of less than five percent. This set of 5kb windows with significant fold change in crossover counts was further filtered to consider only those with six of more crossovers among the fish being compared. Assuming a uniform distribution, the probability of this happening by chance (6 of 18926 crossovers falling within 5kb of each other) is 1.6×10^-30^. Neighboring or overlapping windows were then merged to describe genomic regions containing candidate ecotype-divergent hotpots.

### Inversions

Structural rearrangements such as inversions are common in natural populations and may segregate within the nuclear families used in this study. Parents may be heterozygous or homozygous for DNA sequences that show opposite orientation to the reference genome assembly. In regions where the genomic orientation of the sample is inverted compared to the reference genome and if there is a crossover within that region, there will be false positive crossovers called at both inversion boundaries. Such events will leave a signature distribution of triplet crossovers within a short physical distance. The list of crossovers was examined to find such triplets where the first and the third crossover occurred within 2 Mb physical distance. Across the genome, 8 such regions with multiple offspring having inversion triplets were detected in 7 different chromosomes (Table S7). For these triplets, first and third crossovers were removed and the position of the second crossover was corrected according to the inverted orientation.

### Investigating fitness effects of recombination in a natural hybrid zone

We leveraged the power of a previously described natural marine-freshwater stickleback hybrid zone with large population sizes to explore the potential fitness effects of recombination in the wild (67). DNA from 1045 fish fin-clipped from 8 sites along the River Tyne, East Lothian Scotland (67), was genotyped at 2265 loci across the genome using an Illumina Golden Gate custom genotyping array. Genome-wide genetic ancestry analysis shows that populations from rockpools (site1) and freshwater locations (sites 5-8) at opposite ends of the hybrid zone have pure marine and freshwater genetic ancestry respectively, while populations in the lower reaches of the river (sites 2-4) are comprised of marine, freshwater, hybrid, and recombinant individuals. Adult sticklebacks in the River Tyne die shortly after breeding in the summer resulting in almost completely non-overlapping generations. This provides a valuable opportunity to study cohorts through time - including the differential growth and survival of pure and admixed genotypes.

285 young-of-the-year from the center of the hybrid zone were finclipped, tagged with an elastomer dye, and released, and their DNA genotyped as above. We explored the fitness effects of recombination in the wild in two ways: using standard length as a proxy for fitness (ability to grow), and changes in within-chromosome IV two-locus LD over temporal samples as an indication of differential genotypic survival. Residual standard length reflecting the relative size of the individuals compared the mean of their cohort, was calculated for young-of-the-year sampled from the center of the hybrid zone by removing the effect of sampling date as a factor.

Focusing on chromosome IV, we calculated the minimal number of crossovers needed to explain an individual‘s multilocus genotype at the five above-mentioned adaptive loci. Importantly, this conservative (underestimate) count of crossovers is distinct from a ―hybrid index‖ that is commonly used in hybrid zone studies. Here, individuals carrying a large number of mismatched marine and freshwater genotypes at the five loci will have a high score (maximum possible score for the five loci studied is eight) while marine, freshwater, and F1 hybrids (heterozygous for non-recombined marine and freshwater chromosomes) will have a low score (minimum possible score is 0). We then used linear models to ask whether significant variation in residual standard length can be explained by the number of crossovers among chrIV loci as an independent variable.

### DMC1 antibody production and validation for ChIP sequencing

To produce DMC1 antibody, we expressed stickleback DMC1 as a codon-optimized (GeneArt, Thermo Fisher Scientific) recombinant protein (vector pColdTM TF-DNA, Takara Clontech) to use as the antigen. Following transformation into the expression strain BL21 DE3, a single colony was inoculated in 20 ml LB broth containing 100 μg ml ampicillin for overnight incubation. 10 ml of the overnight culture was inoculated into 500 ml LB without any antibiotic grown to OD 0.4-0.9. Once the culture acquired optimal concentration, it was incubated on ice for 30 minutes and 0.5 mM IPTG was added. Protein expression was carried on overnight at 15^0^C and purified the next day using Ni-NTA columns. The purified recombinant protein was then used as an antigen in an 87 days immunization program on Guinea pigs (outsourced to Eurogentec). The DMC1 antibody was then purified from antisera following affinity purification (using Affi-Gel 15 from Biorad) protocol. Western blot, ChIP, and immunohistochemical staining of histological sections on testes material were used to verify sufficient specificity and affinity.

### ChIP sequencing

Sticklebacks are seasonal breeders and during early winter the testes are enriched for primary spermatocytes undergoing the first round of meiosis (84). In the fish facility, the fish are reared in a cycle of 3 months high light (16 hours light, 8 hours dark), 3 months low light (8 hours light, 16 hours dark). Using Haematoxylin-Eosin staining on histological sections of testes, we determined that in our fish facility, the first round of meiosis in males occurred during the last three weeks of the 3-month low light period. Testes and livers were therefore collected during this time window, snap-frozen in liquid nitrogen, and stored in -80‘ .

For chromatin immunoprecipitation (ChIP) followed by sequencing of proteins of interest, we used a pool of about 20 testes. A pool of about 20 pieces of liver tissue was used as the negative control. A sequential pull down with homemade DMC1 antibody followed by commercial H3K4me3 antibody (Millipore (cat#07-473)) was carried out with the same tissue lysate. DMC1 ChIP and library preparation were carried out following the protocol described in (85, 86). The Kinetic Enrichment of single stranded DNA was excluded for H3K4me3 ChIP. The libraries were sequenced in Illumina Hiseq 3000 with 150 bp paired end cycle. After sequencing, DMC1 ChIP reads were trimmed to 40 bp using Trimmomatic (87), and a specialized bioinformatic pipeline described in Khil et al. 2012 (86) was used for processing the reads. Peaks in the DMC1 ChIP data in comparison with input were called using MACS2 (v2.1.1) (88). with the following parameters:. -q 0.1 --nomodel --slocal 5000 - -llocal 10000 --extsize 800 -f BED --SPMR -g 463000000 -B. For H3K4me3 ChIP reads, macs2 peak calling was done using default parameters.

### ATAC sequencing of primary spermatocytes

Using Haematoxylin-Eosin staining on histological sections of testes (see (84)), we determined that in our fish facility, the first round of meiosis in males occurred during the last three weeks of 3-month low light period. Testes were dissected from adult males during this time window for ATAC-seq and processed immediately after dissection.The isolation of primary spermatocytes by flow cytometry was carried out using a BD FACSMelody Cell Sorter (BD Biosciences): initially, DAPI staining for chromatin content (4C primary spermatocytes, 2C secondary spermatocytes, 1C spermatids/sperm) and microscopy were used to optimize the gating strategy that was eventually based on FSC-A and SSC-A alone (∼ cell size and complexity), since these sufficiently differentiated between the primary spermatocytes, secondary spermatocytes and spermatids/sperm. Briefly, for each ATAC-seq (89) replicate, the testes of three males were pooled, dissociated with 0.07U/ml Liberase (Sigma), 50000 primary spermatocytes were isolated using flow cytometry, cell membranes were lysed with cold 10mM Tris pH8; 10mM NaCl; 3mM MgCl2; 0.1% NP-40 and nuclei were isolated by centrifugation, Tn5 was applied onto the nuclei in standard TAPS-D F buffer for 30min +37‘ with shaking (1000rpm), the tagged DNA was cleaned with the Min Elute kit (Qiagen) and processed to a sequencing library. A control library of naked genomic DNA was also made from each testes pool. The libraries were sequenced on an Illumina platform, the reads mapped to GasAcu1, then mitochondrial reads and PCR duplicates were removed, the files were downsampled to the level of the limiting coverage (20M read pairs) and macs2 was used to call peaks (--shift -100 --extsize 200 --tsize 150). Three marine replicates and three freshwater replicates were analyzed. The replicates of both ecotypes were pooled for comparison to CO coordinates and anti-DMC1-ChIP-seq peaks.

## Acknowledgements

We would like to thank Li Ying Tan, Saad Arif, and Cecilia Martinez for collecting the fishes used in this study and for performing the four field crosses. We are grateful to Frank Chan for input into experimental design, data analysis pipeline, and for all the fruitful discussions. We would like to thank Derek Lundberg and Beth Rowan for their help with developing the home-made multiplexed library preparation protocol. We also thank Christa Lanz and the Genome Center at the Max Planck Institute for Biology, Tübingen for the high throughput sequencing.

## Author contributions

FJ conceived the project and FJ, VV, and EH designed the experiments. VV performed the in vitro crosses the prepared the genomic libraries for whole genome sequencing. VV, AD, and FJ analyzed the whole genome sequencing data and prepared the pipeline for recombination mapping. VV and EH produced the antibody, performed the ChIP sequencing, and analyzed the data. EH performed the ATAC-sequencing and analyzed the data. FJ carried out the hydroid zone field study for which DK contributed towards genotyping array design and SB and DA performed the genotyping. All other data analysis, figure preparation, and manuscript writing was done by VV, EH, AD, and FJ.

