## Supplementary_material for "Fine-scale contemporary recombination variation and its fitness consequences in adaptively diverging stickleback fish"

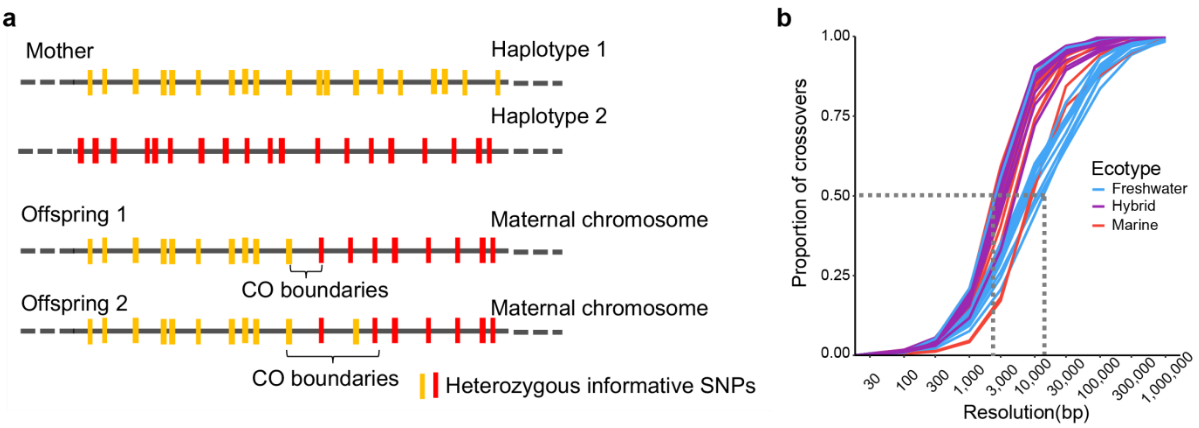


***Fig S1: Crossover definition:*** *a) From phased chromosomes of a parent and offspring, a crossover is defined as an interval flanked by the last SNP of the first haplotype and the first SNP of the second haplotype (eg: offspring 1). In a scenario such as in offspring 2, where two or more switches occurred within the minimum required block size (50kb) of each other, the crossover location was defined by the last SNP of the first large phase block and first SNP of the next large phase block. b) Proportion of crossovers in parents with their respective resolution (interval size within which crossover is identified) is shown. Median crossover resolution is marked with dotted lines.*

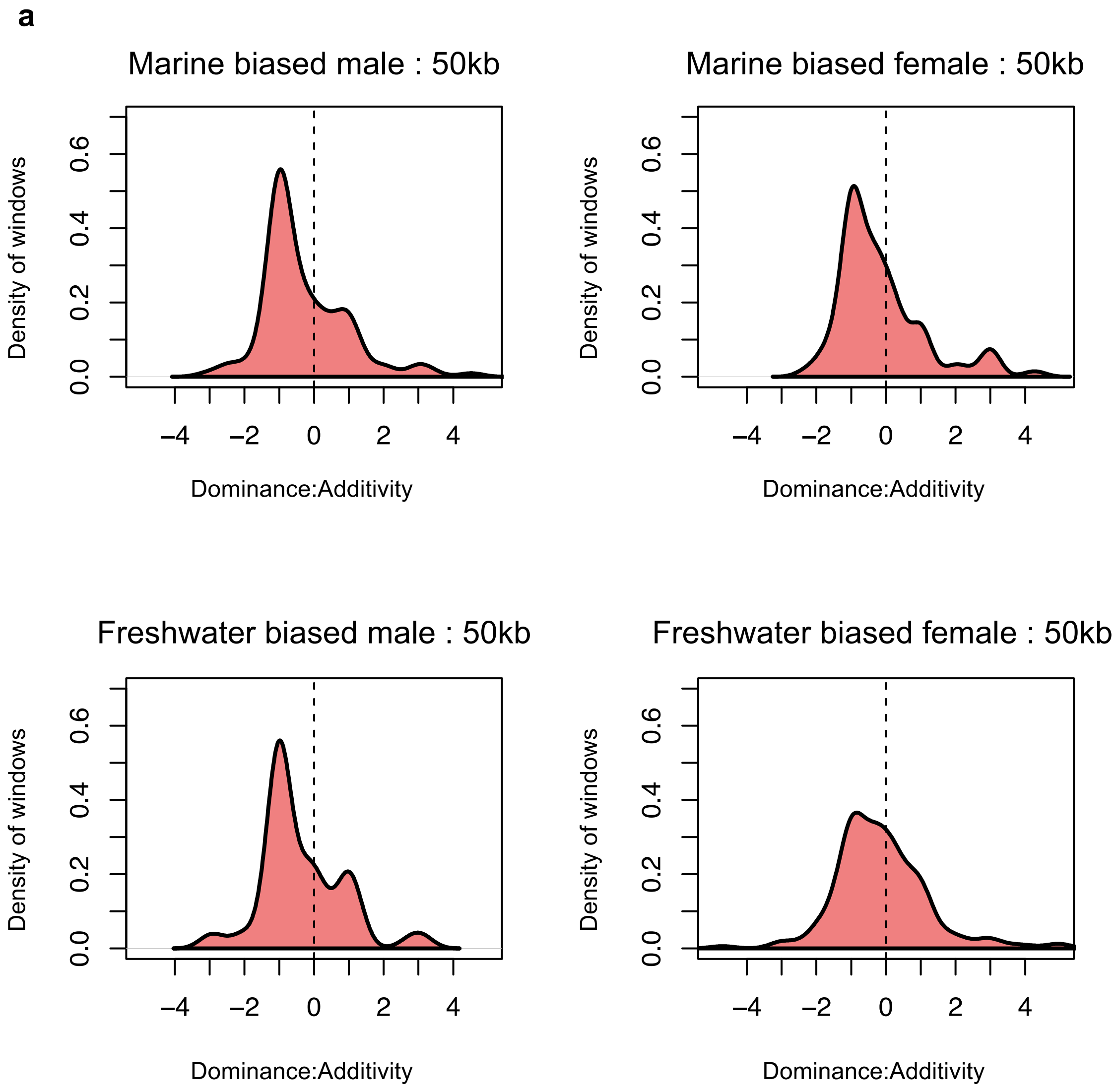


***Fig S2: Partially recessive recombination rates (partial or full dominance of low recombining parental ecotype) in both sexes of hybrids in ecotype diverging windows.*** *a) Density estimates of dominance-additivity ratio of parental ecotype crossover count in hybrid males and females in ecotype diverging windows is shown. A value of zero indicates recombination behaves additively in F1s relative to parents, 1 dominantly, -1 recessively, and values >1 or <-1 imply over or underdominance of recombination respectively. A bias towards low recombining parental ecotype is observed irrespective of which ecotype is paternal and which maternal. Results from analysis at 50kb bresolution are shown. Similar results are obtained with 500kb and 5kb resolutions.*

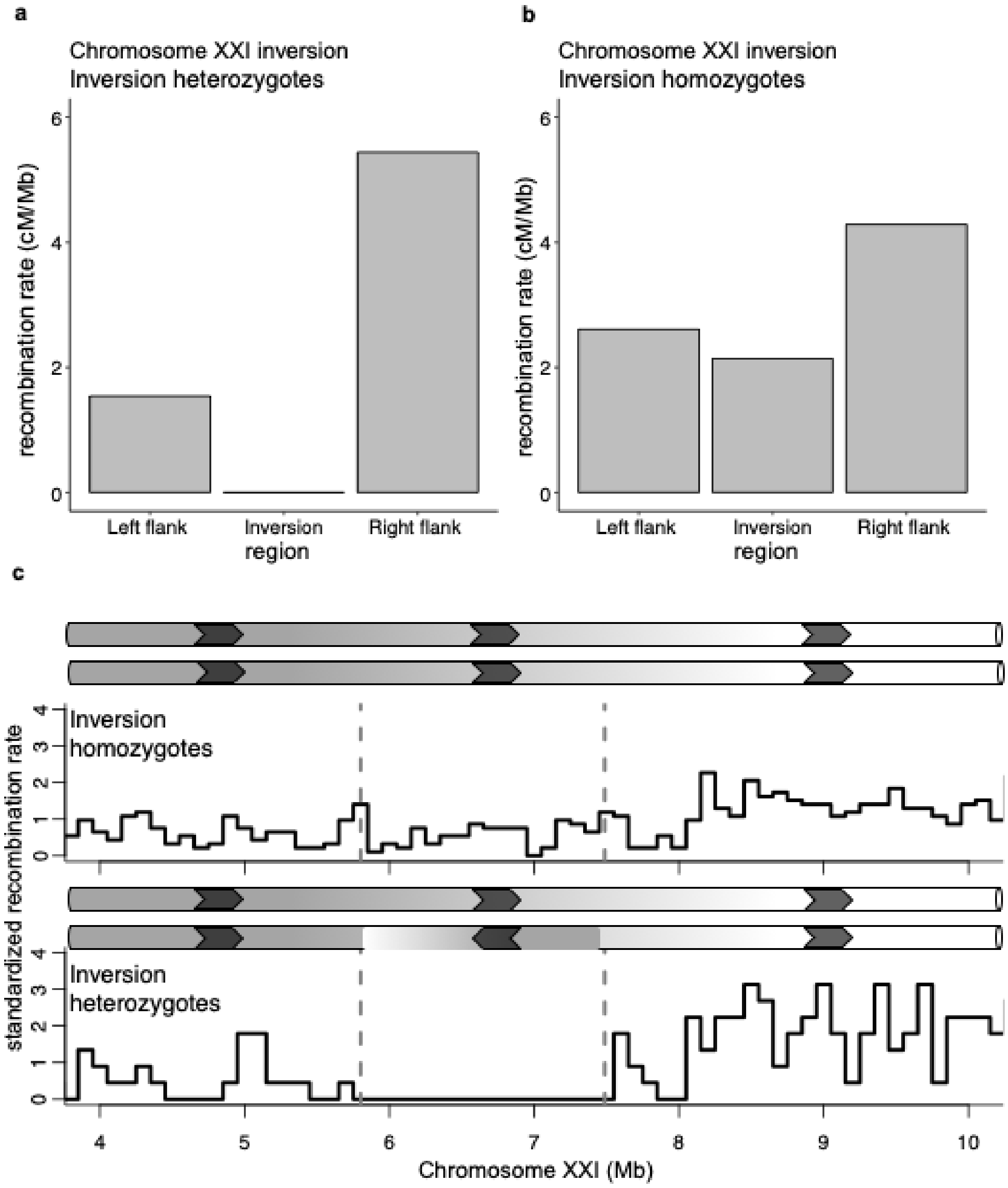


***Fig S3: Crossover suppression within chrXXI inversion heterozygotes.*** *Zero recombination rate within inversion in contrast to left and right flanking regions of the same size is observed in (a) heterozygotes (N=25) but not in (b) homozygotes (N=11). (c)Standardized recombination rate (SRR) in 100 kb sliding windows across a 6 Mb region including the chrXXI inversion. Dotted grey vertical lines mark the inversion boundaries. SRR for inversion homozygotes (top panel) and heterozygotes (bottom panel) are shown with a schematic of the sequence orientation. A complete suppression of recombination within the inversion boundaries is seen in heterozygotes with compensatory increase in recombination on the right flank (closer to telomere).*

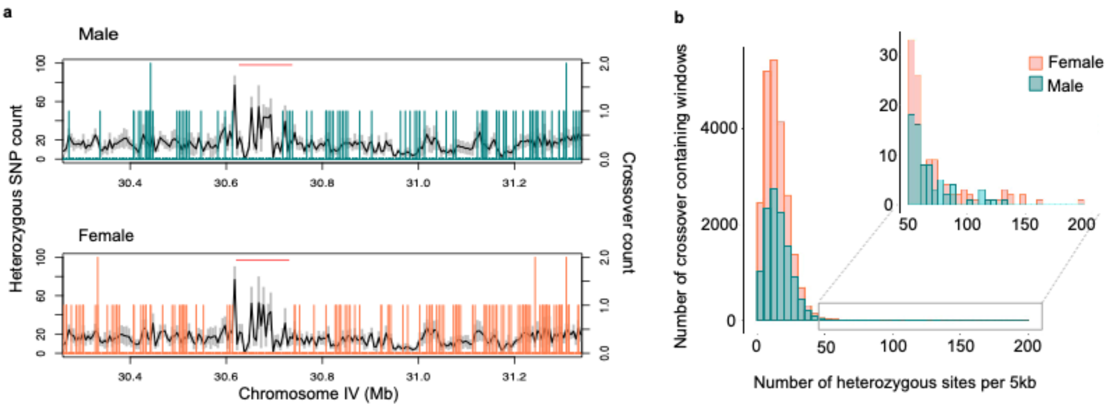


***Fig S4: Crossovers are suppressed in windows with high heterozygosity.*** *a) Recombination suppression is observed in a stretch of high heterozygous windows in an otherwise actively recombining region at the end of chromosome IV. Mean heterozygous SNP counts in 5kb sliding windows (black) along with standard deviation among individuals (grey) are plotted for males (top panel) and females (bottom panel). Crossover counts in corresponding windows are overlaid (green: males, orange: females). Crossover suppressed regions with high heterozygosity are marked with horizontal red lines. b) Histogram of heterozygosity in crossover-containing 5kb windows shows that both male (green) and female (orange) crossovers (COs) mostly occur in windows with less than 50 heterozygous SNP per 5kb. Crossovers are rarely observed in windows with high heterozygosity (zoomed in region).*

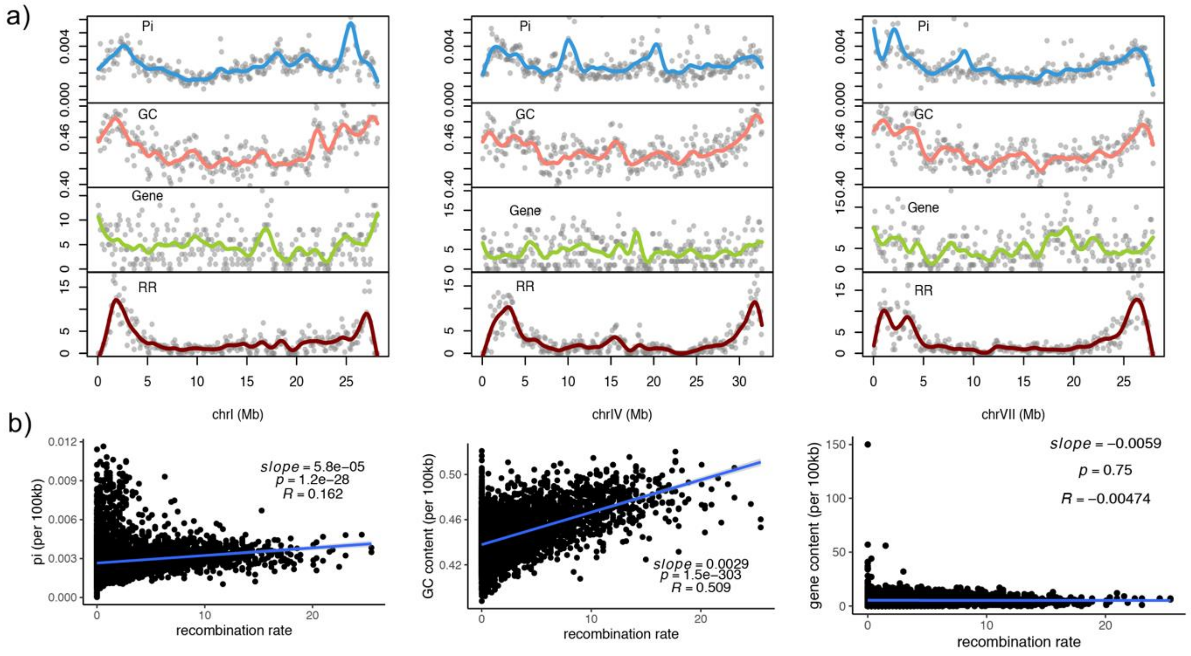


***Fig S5: Recombination rate is positively correlated with GC content and nucleotide diversity but not gene density.*** *a) Recombination rate (RR), GC content, and nucleotide diversity (pi) measured in 100kb sliding windows across three chromosomes (chrI, chrIV, and chrVII) shows a sub-telomeric enrichment. However, gene distribution does not show any obvious pattern and is mostly chromosome specific. b) Linear fit applied on recombination rate versus nucleotide diversity (pi) and GC content across the genome in 100kb windows shows a significant positive correlation, whereas the negative correlation observed between recombination rate and gene density is not statistically significant at this scale.*

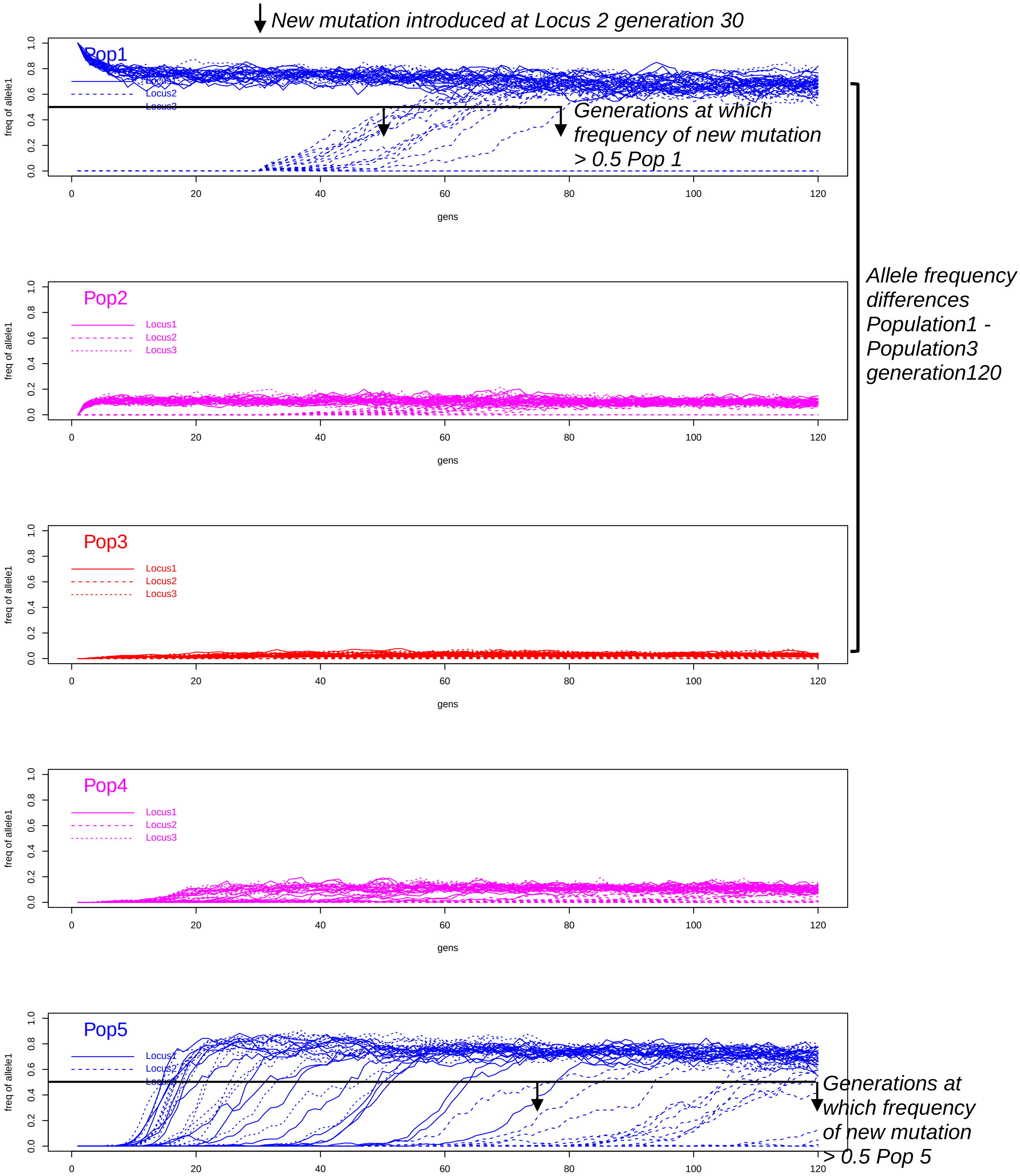


***Fig S6. Forward simulations of three linked loci quantify the rate of the spread of an adaptive allele through hybrid zones maintained in selection migration balance under differing levels of heterochiasmy, recombination suppression, and overall recombination rate. (Simulation1 – No-heterochiasmy, no recombination suppression, medium recombination rate).*** *Divergence between marine and freshwater populations is maintained, despite high rates of gene flow between populations. A new freshwater adaptive mutation is introduced at locus2 in generation 30. Each plot shows the frequency of the freshwater adaptive allele in populations 1 (top) through to 5 (bottom). Colors correspond to the habitats each with different simulated fitness effects of alleles (freshwater – blue; intertidal – magenta; marine – red). See Supplementary Tables S8 and Table S9 and Supplementary methods for more information.*

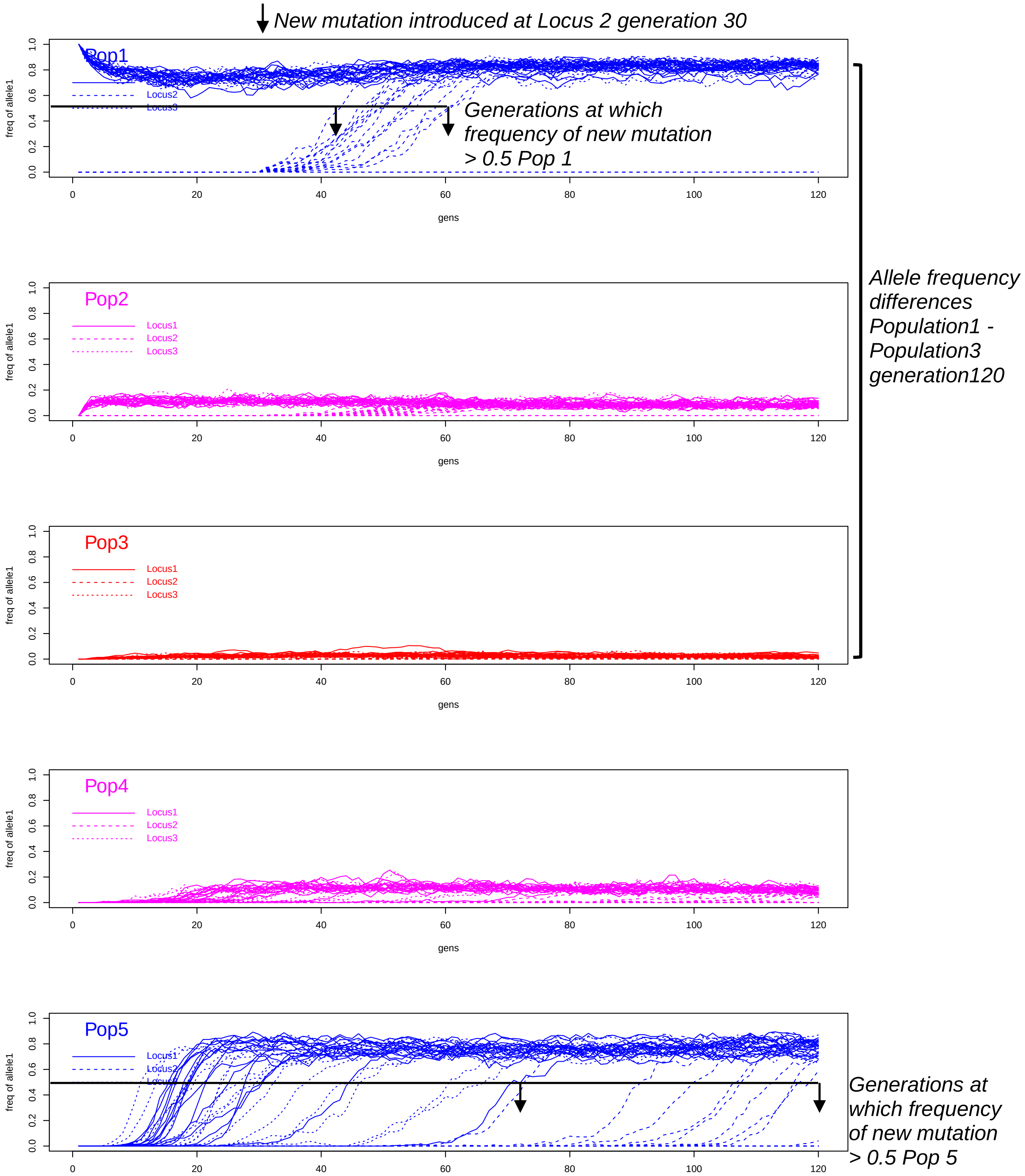


***Fig S7 Forward simulations of three linked loci quantify the rate of the spread of an adaptive allele through hybrid zones maintained in selection migration balance under differing levels of heterochiasmy, recombination suppression, and overall recombination rate. (Simulation2 – No-heterochiasmy, 10fold recombination suppression, medium recombination rate).*** *Divergence between marine and freshwater populations is maintained, despite high rates of gene flow between populations. A new freshwater adaptive mutation is introduced at locus2 in generation 30. Each plot shows the frequency of the freshwater adaptive allele in populations 1 (top) through to 5 (bottom). Colors correspond to the habitats each with different simulated fitness effects of alleles (freshwater – blue; intertidal – magenta; marine – red). See Supplementary Tables S8 and Table S9 and Supplementary methods for more information.*

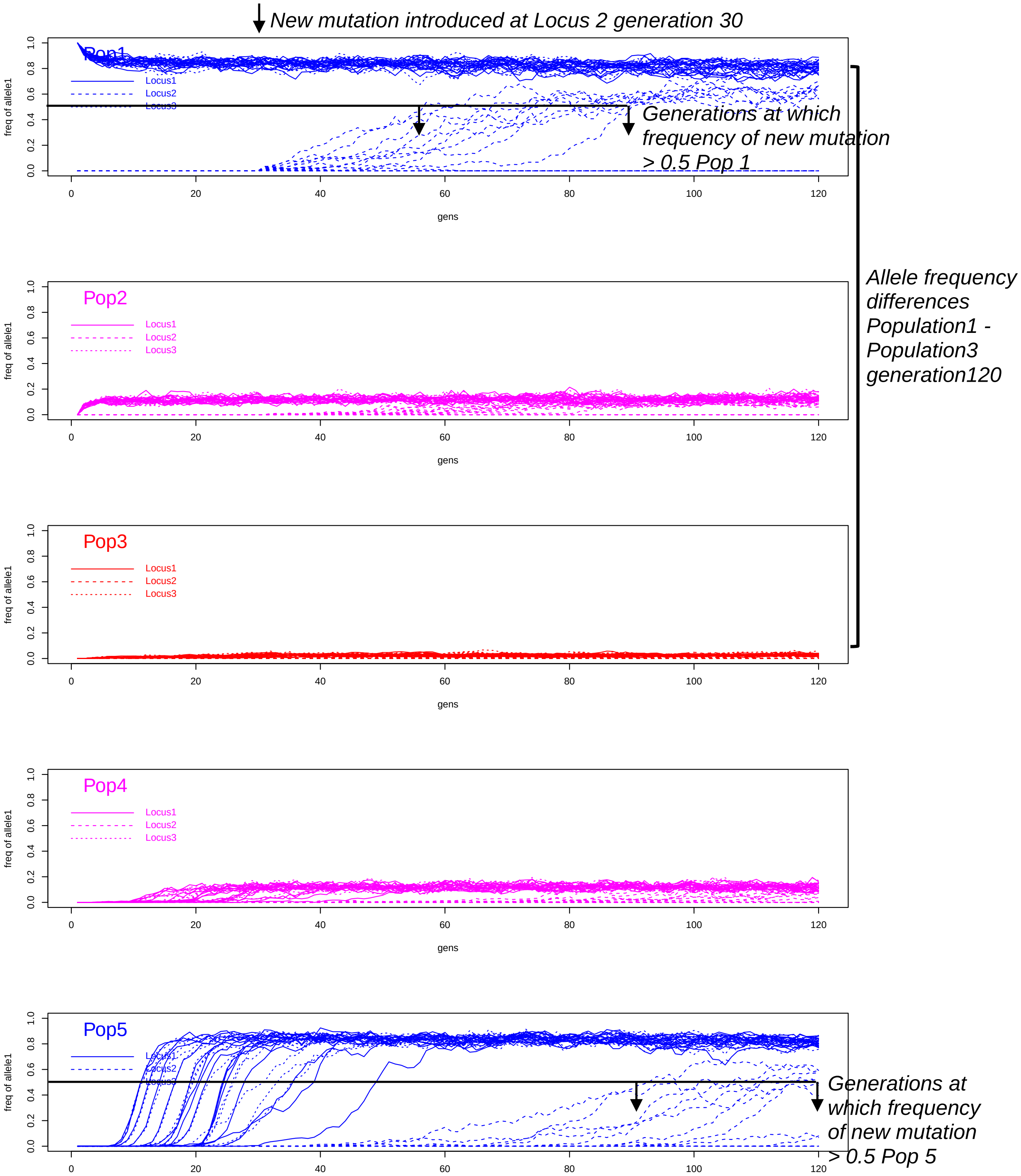


***Fig S8 Forward simulations of three linked loci quantify the rate of the spread of an adaptive allele through hybrid zones maintained in selection migration balance under differing levels of heterochiasmy, recombination suppression, and overall recombination rate. (Simulation3 – Strong heterochiasmy, no recombination suppression, medium recombination rate).*** *Divergence between marine and freshwater populations is maintained, despite high rates of gene flow between populations. A new freshwater adaptive mutation is introduced at locus2 in generation 30. Each plot shows the frequency of the freshwater adaptive allele in populations 1 (top) through to 5 (bottom). Colors correspond to the habitats each with different simulated fitness effects of alleles (freshwater – blue; intertidal – magenta; marine – red). See Supplementary Tables S8 and Table S9 and Supplementary methods for more information.*

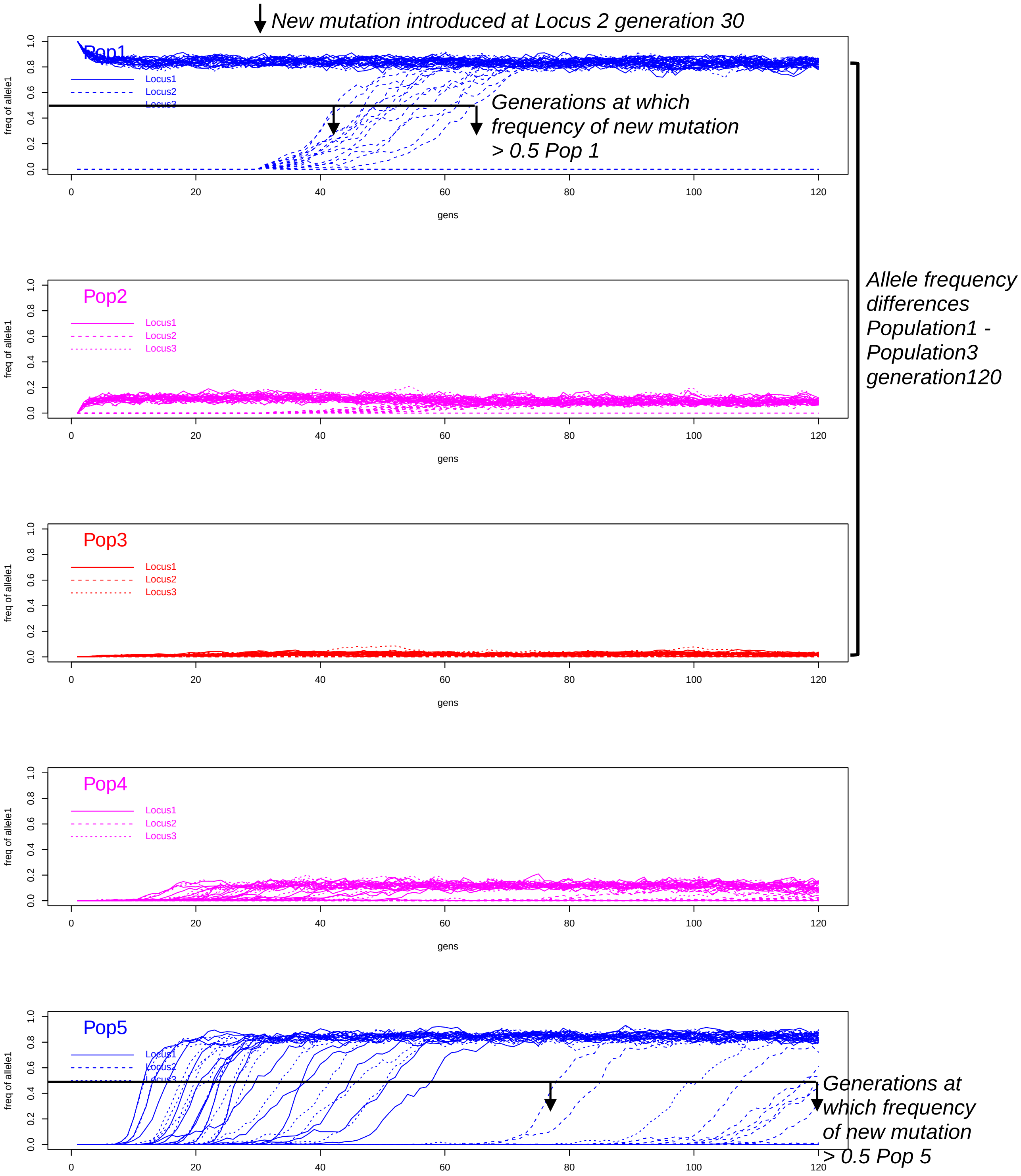


***Fig S9 Forward simulations of three linked loci quantify the rate of the spread of an adaptive allele through hybrid zones maintained in selection migration balance under differing levels of heterochiasmy, recombination suppression, and overall recombination rate. (Simulation4 – Strong heterochiasmy, 10fold recombination suppression, medium recombination rate).*** *Divergence between marine and freshwater populations is maintained, despite high rates of gene flow between populations. A new freshwater adaptive mutation is introduced at locus2 in generation 30. Each plot shows the frequency of the freshwater adaptive allele in populations 1 (top) through to 5 (bottom). Colors correspond to the habitats each with different simulated fitness effects of alleles (freshwater – blue; intertidal – magenta; marine – red). See Supplementary Tables S8 and Table S9 and Supplementary methods for more information.*

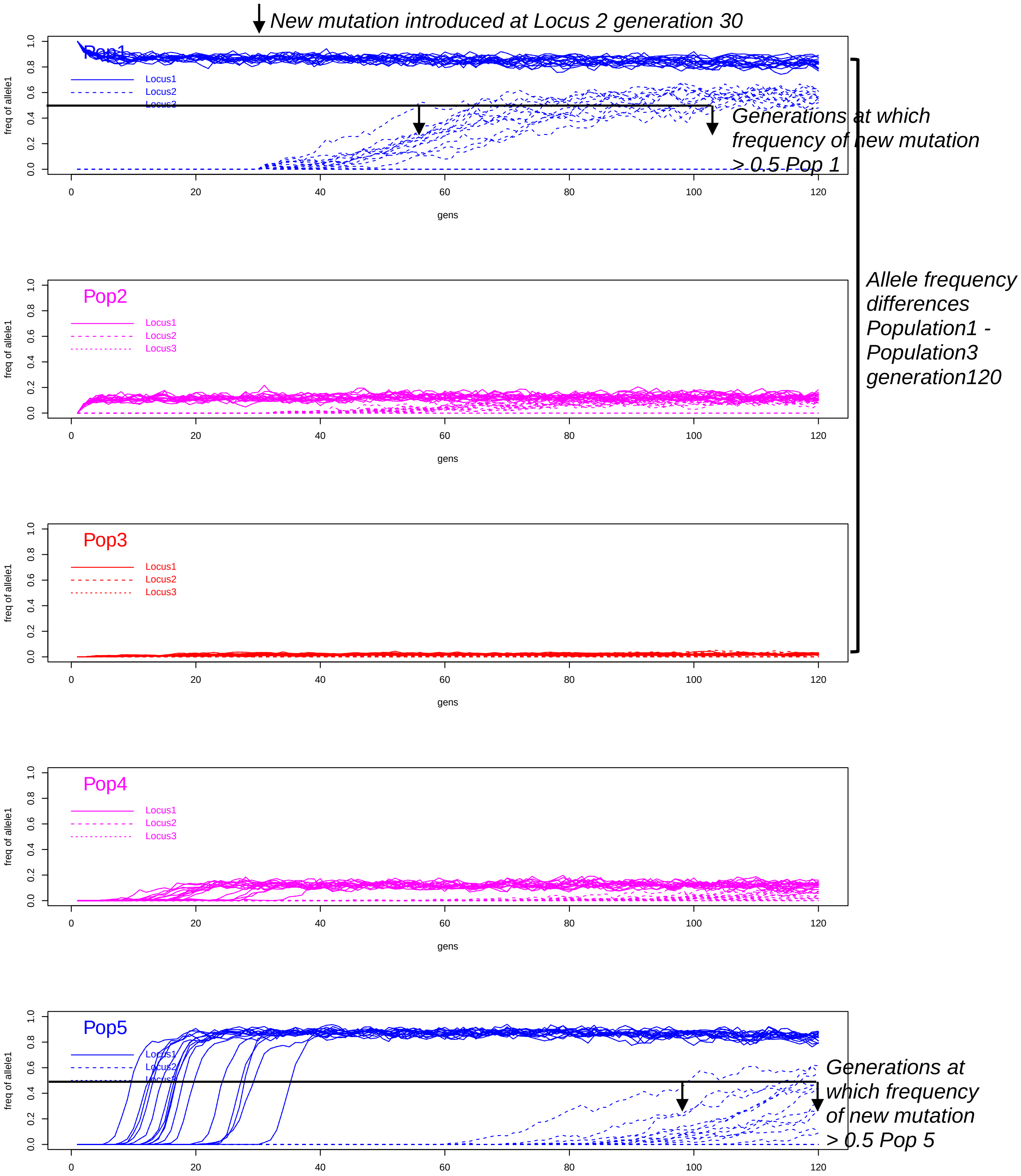


***Fig S10 Forward simulations of three linked loci quantify the rate of the spread of an adaptive allele through hybrid zones maintained in selection migration balance under differing levels of heterochiasmy, recombination suppression, and overall recombination rate. (Simulation5 – No heterochiasmy, no recombination suppression, high recombination rate).*** *Divergence between marine and freshwater populations is maintained, despite high rates of gene flow between populations. A new freshwater adaptive mutation is introduced at locus2 in generation 30. Each plot shows the frequency of the freshwater adaptive allele in populations 1 (top) through to 5 (bottom). Colors correspond to the habitats each with different simulated fitness effects of alleles (freshwater – blue; intertidal – magenta; marine – red). See Supplementary Tables S8 and Table S9 and Supplementary methods for more information.*

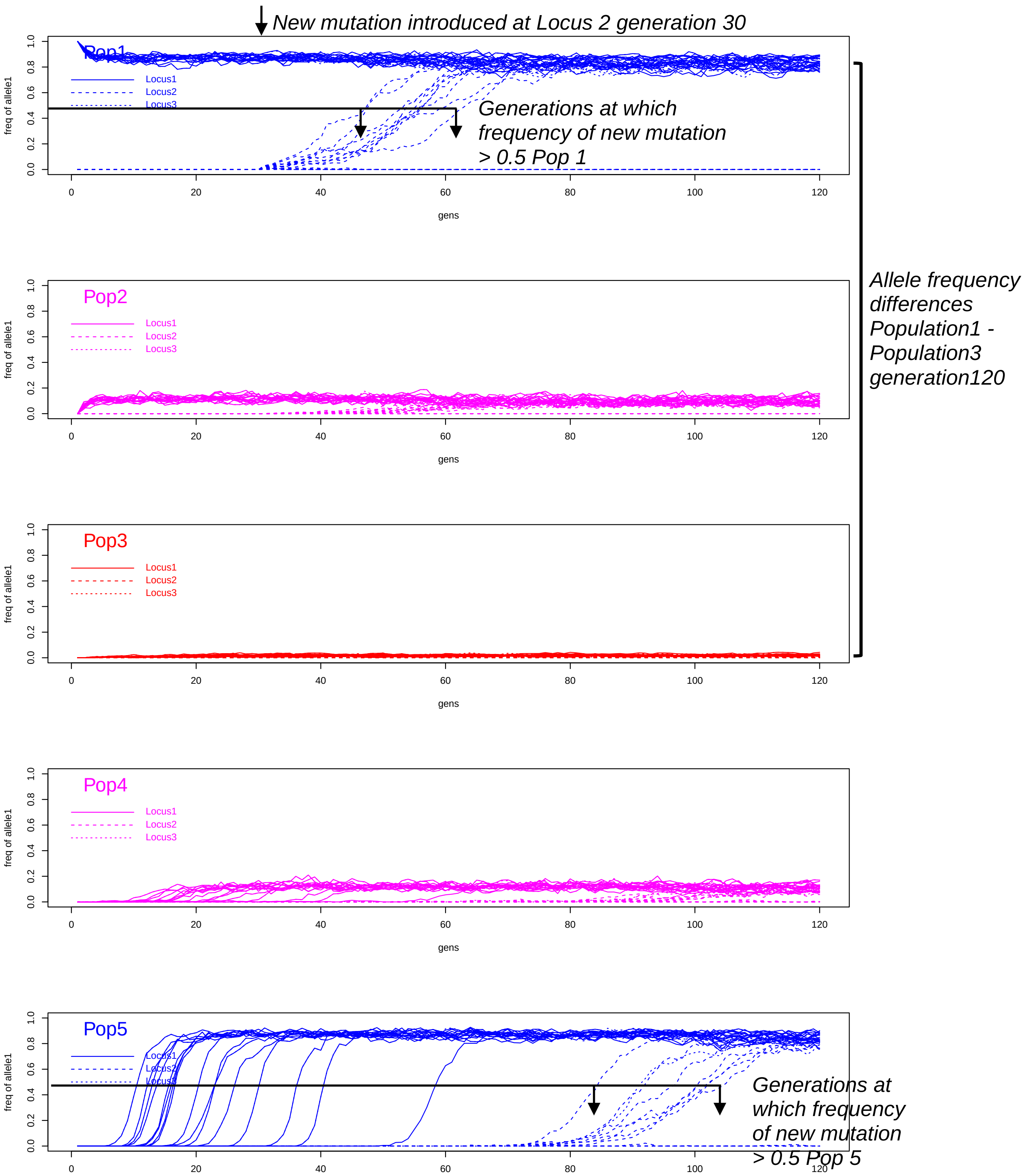


***Fig S11 Forward simulations of three linked loci quantify the rate of the spread of an adaptive allele through hybrid zones maintained in selection migration balance under differing levels of heterochiasmy, recombination suppression, and overall recombination rate. (Simulation6 – No heterochiasmy, 10fold recombination suppression, high recombination rate).*** *Divergence between marine and freshwater populations is maintained, despite high rates of gene flow between populations. A new freshwater adaptive mutation is introduced at locus2 in generation 30. Each plot shows the frequency of the freshwater adaptive allele in populations 1 (top) through to 5 (bottom). Colors correspond to the habitats each with different simulated fitness effects of alleles (freshwater – blue; intertidal – magenta; marine – red). See Supplementary Tables S8 and Table S9 and Supplementary methods for more information.*

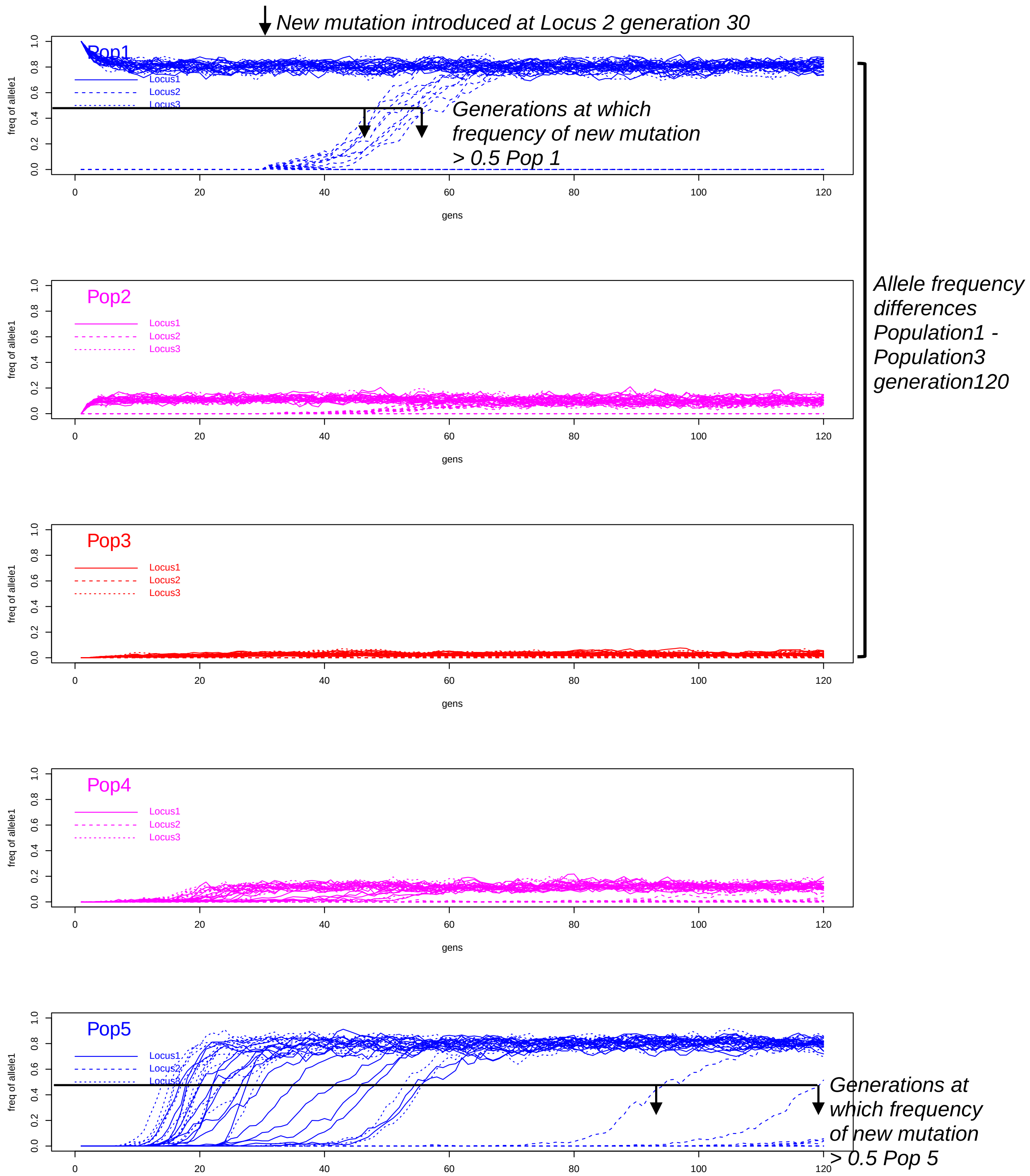


***Fig S12 Forward simulations of three linked loci quantify the rate of the spread of an adaptive allele through hybrid zones maintained in selection migration balance under differing levels of heterochiasmy, recombination suppression, and overall recombination rate. (Simulation7 – No heterochiasmy, no recombination suppression, low recombination rate).*** *Divergence between marine and freshwater populations is maintained, despite high rates of gene flow between populations. A new freshwater adaptive mutation is introduced at locus2 in generation 30. Each plot shows the frequency of the freshwater adaptive allele in populations 1 (top) through to 5 (bottom). Colors correspond to the habitats each with different simulated fitness effects of alleles (freshwater – blue; intertidal – magenta; marine – red). See Supplementary Tables S8, Tables S9 and Supplementary methods for more information.*

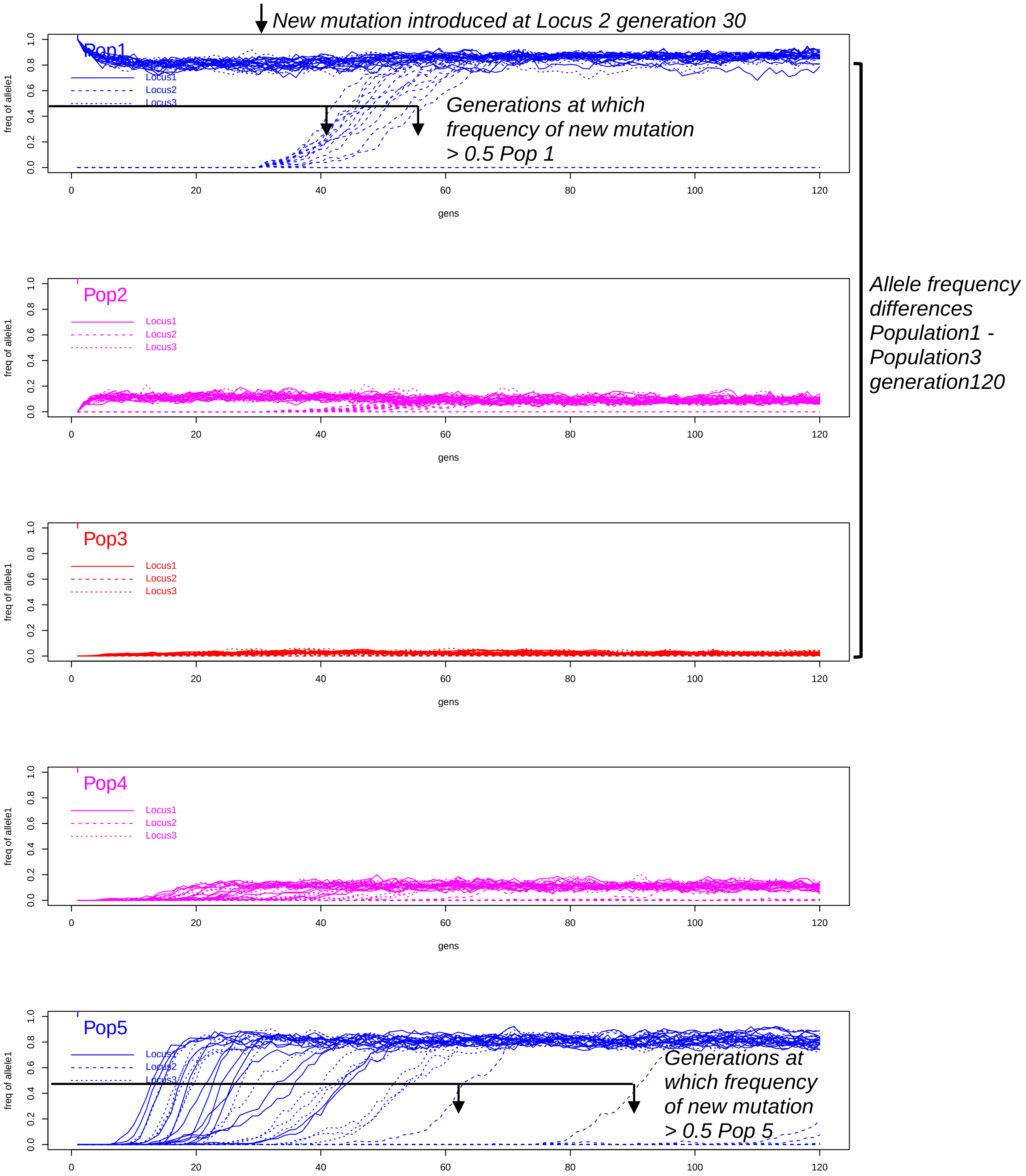


***Fig S13 Forward simulations of three linked loci quantify the rate of the spread of an adaptive allele through hybrid zones maintained in selection migration balance under differing levels of heterochiasmy, recombination suppression, and overall recombination rate. (Simulation8 – No heterochiasmy, 10fold recombination suppression, low recombination rate).*** *Divergence between marine and freshwater populations is maintained, despite high rates of gene flow between populations. A new freshwater adaptive mutation is introduced at locus2 in generation 30. Each plot shows the frequency of the freshwater adaptive allele in populations 1 (top) through to 5 (bottom). Colors correspond to the habitats each with different simulated fitness effects of alleles (freshwater – blue; intertidal – magenta; marine – red). See Supplementary Tables S8 and Table S9 and Supplementary methods for more information.*

| Family | Ecotype | Number of offspring | Dad coverage (x) | Mum coverage (x) | Offspring mean coverage (x) ± SD | Dad informative SNP count | Mum informative SNP count | Dad inter-SNP mean distance (bp) | Mum inter-SNP mean distance (bp) |
|---|---|---|---|---|---|---|---|---|---|
| **X1** | Freshwater | 94 | 38.27 | 49.43 | 13.18 ± 3.50 | 259704 | 250250 | 1541 | 1591 |
| **X4** | Freshwater | 93 | 70.54 | 54.69 | 19.83 ± 6.21 | 385432 | 434153 | 1038 | 922 |
| **X268** | Freshwater | 92 | 236.14 | 188.56 | 14.11 ± 3.60 | 441892 | 282717 | 906 | 1406 |
| **X284** | Freshwater | 91 | 58.65 | 80.06 | 11.86 ± 2.56 | 320492 | 173594 | 1247 | 2301 |
| **X350** | Freshwater | 93 | 60.98 | 67.45 | 13.35 ± 2.44 | 285827 | 238058 | 1401 | 1679 |
| **X351** | Freshwater | 93 | 229.45 | 298.05 | 13.38 ± 2.68 | 350647 | 283794 | 1141 | 1410 |
| **X11** | Marine | 94 | 77.81 | 63.65 | 11.65 ± 2.17 | 385709 | 319840 | 1039 | 1252 |
| **X20** | Marine | 91 | 57.69 | 138.48 | 14.07 ± 5.48 | 325633 | 307571 | 1230 | 1300 |
| **X291** | Marine | 94 | 57.58 | 70.09 | 12.46 ± 2.97 | 465765 | 364930 | 860 | 1098 |
| **X294** | Marine | 94 | 119.55 | 117.69 | 14.11 ± 2.99 | 492870 | 410454 | 813 | 976 |
| **X295** | Marine | 93 | 35.04 | 44.24 | 13.24 ± 3.60 | 216217 | 111255 | 1851 | 3586 |
| **X296** | Marine | 93 | 56.15 | 68.44 | 13.62 ± 2.89 | 465613 | 246504 | 861 | 1625 |
| **X273** | Hybrid | 94 | 62.47 | 64.30 | 13.45 ± 2.71 | 461511 | 345952 | 868 | 1155 |
| **X274** | Hybrid | 94 | 87.67 | 72.32 | 13.63 ± 2.99 | 451754 | 312601 | 887 | 1282 |
| **X800** | Hybrid | 94 | 32.82 | 45.05 | 13.32 ± 2.84 | 269782 | 323189 | 1477 | 1239 |
| **X366** | Hybrid | 86 | 69.86 | 59.70 | 13.36 ± 3.68 | 474780 | 329376 | 844 | 1216 |
| **X389** | Hybrid | 93 | 62.58 | 69.91 | 14.63 ± 2.35 | 513093 | 385061 | 781 | 1041 |
| **X391** | Hybrid | 93 | 63.70 | 57.80 | 14.57 ± 2.68 | 501055 | 367392 | 798 | 1090 |

***Table S1 : Family, sequencing, SNP summary table***

| merged divergent hotspot region | chr | start | stop | Marine CO Count[1] | Freshwater CO Count[1] | Hybrid CO Count[1] | Direction of Divergence in CO Counts |
|---|---|---|---|---|---|---|---|
| *chrXII:1333868-1339417* | chrXII | 1333868 | 1339417 | 3 | 8 | 2 | Freshwater>Hybrid |
| chrXVIII:2579793-2584793 | chrXVIII | 2579793 | 2584793 | 1 | 5 | 3 | Freshwater>Marine |
| chrXVIII:14592056-14597056 | chrXVIII | 14592056 | 14597056 | 1 | 5 | 4 | Freshwater>Marine |
| chrX:13823856-13828856 | chrX | 13823856 | 13828856 | 0 | 6 | 1 | Freshwater>Marine; Freshwater>Hybrid |
| chrXIII:18523940-18529348 | chrXIII | 18523940 | 18529348 | 1 | 7 | 2 | Freshwater>Marine; Freshwater>Hybrid |
| *chrXXI:11393230-11406225* | chrXXI | 11393230 | 11406225 | 9 | 17 | 5 | Freshwater>Marine; Freshwater>Hybrid |
| *chrI:1543413-1554456* | chrI | 1543413 | 1554456 | 7 | 5 | 3 | Marine>Freshwater; Freshwater>Marine; Marine>Hybrid [a] |
| chrIII:16405102-16412742 | chrIII | 16405102 | 16412742 | 5 | 1 | 7 | Hybrid>Freshwater |
| chrVII:2397086-2404154 | chrVII | 2397086 | 2404154 | 0 | 1 | 7 | Hybrid>Marine |
| chrXIX:1449114-1455016 | chrXIX | 1449114 | 1455016 | 2 | 2 | 6 | Hybrid>Marine |
| chrXI:14097150-14102150 | chrXI | 14097150 | 14102150 | 7 | 1 | 3 | Marine>Freshwater; Marine>Hybrid |
| chrXIX:1623924-1631003 | chrXIX | 1623924 | 1631003 | 10 | 4 | 3 | Marine>Hybrid |

1 Counts of all crossover midpoints (not downsampled, but excluding crossovers that overlap scaffold boundaries and/or have interval resolution greater than 10kb) that fall within the merged genomic region are shown for each ecotype. a Neighbouring 5kb windows within this region show crossover counts with freshwater>marine, marine>freshwater, and marine>hybrids

***Table S2:*** *Genomic regions containing 5kb hotspot windows with >=6 total crossovers and significant differences in crossover counts between marine, freshwater and hybrid fish. Statistical tests were performed in 5kb intervals on equal numbers of crossovers from all three ecotypes (see Tables S3-S5), and merged to generate intervals shown in this table. Assuming a uniform distribution, the probability for 6 of 18926 crossovers falling within 5kb of each other by chance is* $1.6 \times 10^{-30}$.

| Freshwater Crossover 5kb Window [1] | chr | start | stop | Freshwater Crossover Count | Marine Crossover Count | Hybrid Crossover Count | log2FoldChange [Marine Crossover Count / Freshwater Crossover Count] [2] | log2FoldChange [Hybrid Crossover Count / Freshwater Crossover Count] [2] | significant_contrast |
|---|---|---|---|---|---|---|---|---|---|
| chrI:1547951-1552951 | chrI | 1547951 | 1552951 | 1 | 5 | 1 | **2.21299** | 0 | Freshwater_v_Marine |
| chrIII:16407356-16412356 | chrIII | 16407356 | 16412356 | 1 | 3 | 6 | 1.49476 | **2.47131** | Freshwater_v_Hybrid |
| chrX:13823856-13828856 | chrX | 13823856 | 13828856 | 6 | 0 | 0 | **-5.93074** | **-5.93074** | Freshwater_v_Marine; Freshwater_v_Hybrid |
| chrXII:1333868-1338868 | chrXII | 1333868 | 1338868 | 7 | 3 | 0 | -1.19555 | **-6.14975** | Freshwater_v_Hybrid |
| chrXII:1333976-1338976 | chrXII | 1333976 | 1338976 | 6 | 3 | 0 | -0.976541 | **-5.93074** | Freshwater_v_Hybrid |
| chrXII:1334417-1339417 | chrXII | 1334417 | 1339417 | 6 | 2 | 0 | -1.53842 | **-5.93074** | Freshwater_v_Hybrid |
| chrXIII:18523940-18528940 | chrXIII | 18523940 | 18528940 | 6 | 1 | 2 | **-2.47131** | -1.53842 | Freshwater_v_Marine |
| chrXIII:18524348-18529348 | chrXIII | 18524348 | 18529348 | 6 | 1 | 1 | **-2.47131** | **-2.47131** | Freshwater_v_Marine; Freshwater_v_Hybrid |
| chrXXI:11393230-11398230 | chrXXI | 11393230 | 11398230 | 9 | 4 | 2 | -1.15024 | **-2.11548** | Freshwater_v_Hybrid |
| chrXXI:11397018-11402018 | chrXXI | 11397018 | 11402018 | 6 | 1 | 3 | **-2.47131** | -0.976541 | Freshwater_v_Marine |
| chrXXI:11400304-11405304 | chrXXI | 11400304 | 11405304 | 8 | 3 | 0 | -1.38565 | **-6.33985** | Freshwater_v_Hybrid |
| chrXXI:11401020-11406020 | chrXXI | 11401020 | 11406020 | 7 | 3 | 0 | -1.19555 | **-6.14975** | Freshwater_v_Hybrid |
| chrXXI:11401225-11406225 | chrXXI | 11401225 | 11406225 | 6 | 3 | 0 | -0.976541 | **-5.93074** | Freshwater_v_Hybrid |

1 Freshwater 5kb crossover hotspot windows with significant fold change difference in crossover counts relative to marine and hybrid ecotypes (FDR <0.05) and >=6 total crossovers in the focal window. Assuming a uniform distribution, the probability of this happening by chance (6 of 18926 crossovers falling within 5kb of each other) is 1.6x10^-30.
2 log2FoldChange q<0.05 shown in bold. Pseudocount of 0.1 added to both sides of log ratio.

***Table S3:*** *5kb windows with significant differences in crossover counts between freshwater and marine fish and freshwater and hybrid fish. For each focal window with >=6 total crossovers among the ecotypes being compared, down-sampled crossover counts for all three ecotypes are shown with fold change ratios of qvalue <0.05 in bold. We included only crossovers not spanning scaffold boundaries, and with crossover interval resolution <=10000bp, down-sampled to generate equal numbers of crossovers per ecotype.*

| Marine Crossover 5kb Window [1] | chr | start | stop | Marine CO Count | Freshwater CO Count | Hybrid CO Count | log2FoldChange [Freshwater CO Count / Marine CO Count] [2] | log2FoldChange [Hybrid CO Count / Marine CO Count] [2] | significant_contrast |
|---|---|---|---|---|---|---|---|---|---|
| chrI:1543413-1548413 | chrI | 1543413 | 1548413 | 1 | 5 | 2 | **2.21299** | 0.932886 | Marine_v_Freshwater |
| chrXI:14097150-14102150 | chrXI | 14097150 | 14102150 | 6 | 1 | 1 | **-2.47131** | **-2.47131** | Marine_v_Freshwater; Marine_v_Hybrid |
| chrXIX:1449114-1454114 | chrXIX | 1449114 | 1454114 | 2 | 2 | 6 | 0 | **1.53842** | Marine_v_Hybrid |
| chrXIX:1624328-1629328 | chrXIX | 1624328 | 1629328 | 8 | 4 | 1 | -0.982298 | **-2.88042** | Marine_v_Hybrid |
| chrXIX:1625740-1630740 | chrXIX | 1625740 | 1630740 | 8 | 3 | 0 | -1.38565 | **-6.33985** | Marine_v_Hybrid |
| chrXIX:1626003-1631003 | chrXIX | 1626003 | 1631003 | 7 | 3 | 0 | -1.19555 | **-6.14975** | Marine_v_Hybrid |
| chrXVIII:2579793-2584793 | chrXVIII | 2579793 | 2584793 | 1 | 5 | 2 | **2.21299** | 0.932886 | Marine_v_Freshwater |
| chrXVIII:14592056-14597056 | chrXVIII | 14592056 | 14597056 | 1 | 5 | 2 | **2.21299** | 0.932886 | Marine_v_Freshwater |

1 Marine 5kb crossover hotspot windows with significant fold change difference in crossover counts relative to freshwater and hybrid ecotypes (FDR <0.05) and >=6 total crossovers in the focal window. Assuming a uniform distribution, the probability of this happening by chance (6 of 18926 crossovers falling within 5kb of each other) is 1.6x10^-30.
2 log2FoldChange q<0.05 shown in bold

***Table S4:*** *5kb windows with significant differences in crossover counts between marine and freshwater fish and marine and hybrid fish. For each focal window with >=6 total crossovers among the ecotypes being compared, down-sampled crossover counts for all three ecotypes are shown with fold change ratios of qvalue <0.05 in bold. We included only crossovers not spanning scaffold boundaries, and with crossover interval resolution <=10000bp, down-sampled to generate equal numbers of crossovers per ecotype.*

| Hybrid Crossover 5kb Window [1] | chr | start | stop | Hybrid CO Count | Marine CO Count | Freshwater CO Count | log2FoldChange [Marine CO Count / Hybrid CO Count] [2] | log2FoldChange [Freshwater CO Count / Hybrid CO Count] [2] | significant_contrast |
|---|---|---|---|---|---|---|---|---|---|
| chrI:1549456-1554456 | chrI | 1549456 | 1554456 | 1 | 5 | 0 | **2.21299** | -3.45943 | Hybrid_v_Marine |
| chrIII:16405102-16410102 | chrIII | 16405102 | 16410102 | 7 | 4 | 1 | -0.792195 | **-2.69032** | Hybrid_v_Freshwater |
| chrIII:16407742-16412742 | chrIII | 16407742 | 16412742 | 6 | 2 | 0 | -1.53842 | **-5.93074** | Hybrid_v_Freshwater |
| chrVII:2397086-2402086 | chrVII | 2397086 | 2402086 | 6 | 0 | 1 | **-5.93074** | -2.47131 | Hybrid_v_Marine |
| chrVII:2399154-2404154 | chrVII | 2399154 | 2404154 | 6 | 0 | 1 | **-5.93074** | -2.47131 | Hybrid_v_Marine |
| chrXIX:1450016-1455016 | chrXIX | 1450016 | 1455016 | 6 | 1 | 2 | **-2.47131** | -1.53842 | Hybrid_v_Marine |
| chrXIX:1623924-1628924 | chrXIX | 1623924 | 1628924 | 2 | 8 | 4 | **1.94753** | 0.965235 | Hybrid_v_Marine |
| chrXIX:1625364-1630364 | chrXIX | 1625364 | 1630364 | 1 | 7 | 3 | **2.69032** | 1.49476 | Hybrid_v_Marine |
| chrXXI:11400166-11405166 | chrXXI | 11400166 | 11405166 | 2 | 3 | 8 | 0.561879 | **1.94753** | Hybrid_v_Freshwater |
| chrXXI:11400197-11405197 | chrXXI | 11400197 | 11405197 | 1 | 3 | 8 | 1.49476 | **2.88042** | Hybrid_v_Freshwater |

1 Hybrid 5kb crossover hotspot windows with significant fold change difference in crossover counts relative to freshwater and marine ecotypes (FDR <0.05) and >=6 total crossovers in the focal window. Assuming a uniform distribution, the probability of this happening by chance (6 of 18926 crossovers falling within 5kb of each other) is 1.6x10^-30.
2 log2FoldChange q<0.05 shown in bold

***Table S5:*** *5kb windows with significant differences in crossover counts between hybrid and freshwater fish and hybrid and marine fish. For each focal window with >=6 total crossovers among the ecotypes being compared, dow- sampled crossover counts for all three ecotypes are shown with fold change ratios of qvalue <0.05 shown in bold. We included only crossovers not spanning scaffold boundaries, and with crossover interval resolution <=10000bp, down-sampled to generate equal numbers of crossovers per ecotype.*

| Male | | | Female | | |
|---|---|---|---|---|---|
| Motif | Probability | Frequency at CO sites | Motif | Probability | Frequency at CO sites |
| 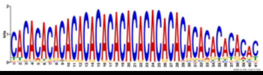 | 3.1e-090 | 25.5% | 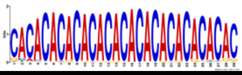 | 2.4e-139 | 23.2% |
| 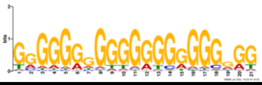 | 9.4e-041 | 51.3% | 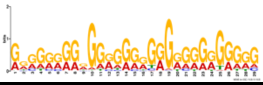 | 6.8e-028 | 39.2% |
| 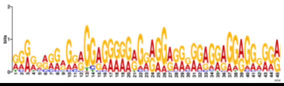 | 2.3e-027 | 1.6% | 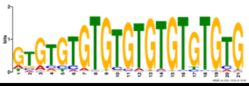 | 5.0e-019 | 33.6% |

***Table S6** : Motifs discovered using MEME considering all male and female high-resolution (<5kb) crossover intervals.*

| Sex | Inversion at | Inversion detected families | Ecotype |
|---|---|---|---|
| **Male** | chrI: 25,264,236 - 25,720,158 | X294 | Marine |
| | chrII: 22,372,205 - 23,174,871 | X268, X294, X296, X350, X351, X366, X389, X391 | Marine, FW, Hybrid B |
| | chrIX: 5,655,297 - 7,950,866 | X389 | Hybrid B |
| | chrXI: 5,431,984 - 5,868,073 | X291 | Marine |
| | chrXI: 15,730,574 - 16,638,140 | X11, X20, X268, X284, X291, X294, X296, X366, X391 | Marine, FW, Hybrid B |
| | chrXVI: 17,195,676 - 17,968,133 | X11, X20, X268, X273, X274, X284, X294, X296, X350, X351, X366, X389, X4, X800 | Marine, FW, Hybrid B, Hybrid A |
| | chrXVII: 641211 - 769373 | X391 | Hybrid B |
| | chrXXI : 5,681,441 - 7,787,895 | X1, X11, X284, X294, X296, X350, X351, X366, X389, X4, X800 | Marine, FW, Hybrid B, Hybrid A |
| **Female** | chrI: 25,267,445 - 25,725,739 | X11, X291 | Marine |
| | chrII: 22,358,302 - 23,062,538 | X1, X268, X366, X800 | Marine, FW, Hybrid A, Hybrid B |
| | chrIX : 5,647,525 - 7,282,910 | X11, X366 | Marine, Hybrid B |
| | chrXI : 5431082 - 5858743 | X11, X20, X294, X389, X800 | Marine, Hybrid A, Hybrid B |
| | chrXI: 15,734,076 - 16,500,546 | X1, X11, X273, X296, X800 | Marine, FW, Hybrid A, |
| | chrXVI: 15,870,586 - 17,836,051 | X1, X11, X273, X20, X274, X284, X291, X294, X296, X350, X351, X366, X389, X391, X4, X800 | Marine, FW, Hybrid A, Hybrid B |
| | chrXVII: 641353 - 769480 | X350 | FW |
| | chrXXI: 5,726,992 -7,494,035 | X11, X268, X273, X284, X291, X294, X295, X296,<br>X350,351, X366, X389, X391, X800 | Marine, FW, Hybrid A, Hybrid B |
| ***Hybrid A: (FW x Marine)F1; Hybrid B: (Mar x FW)F1** | | | |

***Table S7:*** *Details of chromosomal inversions (compared to reference genome) detected in this data set.*

| simulation | recombination rate female (cM) | recombination rate male (cM) | effective sex-averaged recombination rate | recombination suppression in cis(coupled)-heterozygotes | Population 1: mean+/-SD generation to reach frequency 0.5 | Population 1: probability of reaching frequency 0.5 within 120 generations | Population 5: mean+/-SD generation to reach frequency 0.5 | Population 5: probability of reaching frequency 0.5 within 120 generations | Locus 1: Mean allele frequency difference at generation 120 between freshwater population 1 and marine population 3 (Δp +/- SD) | Locus 2: Mean allele frequency difference at generation 120 between freshwater population 1 and marine population 3 (Δp +/- SD) | Locus 3: Mean allele frequency difference at generation 120 between freshwater population 1 and marine population 3 (Δp +/- SD) | Figure | Description |
|---|---|---|---|---|---|---|---|---|---|---|---|---|---|
| 1 | 0.55 | 0.55 | 0.55 | none | 62.7+/-7.4 | 0.8 | 102.7+/-13.9 | 0.6 | 0.66+/-0.06 | 0.49+/-0.25 | 0.65+/-0.06 | Supp Fig S6 | No-heterochiasmy, No recombination suppression, medium recombination rate |
| 2 | 0.55 | 0.55 | 0.55 | 10fold | 52.2+/-5.4 | 0.9 | 103.4+/-14.9 | 0.5 | 0.80+/-0.03 | 0.73+/-0.29 | 0.80+/-0.04 | Supp Fig S7 | No-heterochiasmy, 10fold recombination suppression, medium recombination rate |
| 3 | 1 | 0.1 | 0.55 | none | 74.4+/-11.1 | 0.6 | 107.5+/-9.1 | 0.4 | 0.79+/-0.04 | 0.33+/-0.30 | 0.78+/-0.04 | Supp Fig S8 | Strong-heterochiasmy, No recombination suppression, medium recombination rate |
| 4 | 1 | 0.1 | 0.55 | 10fold | 53.2+/-7.0 | 0.8 | 101.8+/-18.8 | 0.3 | 0.81+/-0.02 | 0.68+/-0.34 | 0.81+/-0.02 | Supp Fig S9 | Strong-heterochiasmy, 10fold recombination suppression, medium recombination rate |
| 5 | 1 | 1 | 1 | none | 79.6+/-12.9 | 0.8 | 111.0+/-9.6 | 0.2 | 0.81+/-0.04 | 0.41+/-0.25 | 0.81+/-0.04 | Supp Fig S10 | No-heterochiasmy, No recombination suppression, high recombination rate |
| 6 | 1 | 1 | 1 | 10fold | 55.7+/-5.1 | 0.6 | 98.4+/-7.3 | 0.4 | 0.80+/-0.05 | 0.51+/-0.40 | 0.82+/-0.03 | Supp Fig S11 | No-heterochiasmy, 10fold recombination suppression, high recombination rate |
| 7 | 0.1 | 0.1 | 0.1 | none | 53.6+/-3.7 | 0.6 | 107.5+/-17.7 | 0.1 | 0.77+/-0.05 | 0.47+/-0.43 | 0.78+/-0.03 | Supp Fig S12 | No-heterochiasmy, No recombination suppression, low recombination rate |
| 8 | 0.1 | 0.1 | 0.1 | 10fold | 48.8+/-4.2 | 0.6 | 78.0+/-19.8 | 0.1 | 0.84+/-0.04 | 0.65+/-0.39 | 0.83+/-0.04 | Supp Fig S13 | No-heterochiasmy, 10fold recombination suppression, low recombination rate |

***Table S8**: Forward simulations quantify the rate of the spread of an adaptive allele through hybrid zones maintained in selection migration balance under differing levels of heterochiasmy, recombination suppression, and overall recombination rate. Summary statistics quantify the generation in which the adaptive allele frequency at Locus 2 reaches a frequency greater than 0.5 in Population1 and 5, the probability of this occurring within 120 generations (proportion of occurrences in all replicate runs), and the mean allele frequency difference +/- standard deviation that is maintained between freshwater population 1 and marine population 3 at each of the three loci at generation 120. Figure numbers corresponding to plots of allele frequency changes over time in each of the populations are shown for each of the simulation conditions in column 12.*

**2-locus haplotype**

| L1 L3 | Marine | Intertidal | Freshwater |
|---|---|---|---|
| M M | 0.5 | 0.5 | 0.05 |
| F M | 0.225 | 0.25 | 0.225 |
| M F | 0.225 | 0.25 | 0.225 |
| F F | 0.05 | 0.5 | 0.5 |

**3-locus haplotype**

| L1L2L3 | Marine | Intertidal | Freshwater |
|---|---|---|---|
| MMM | 0.5 | 0.25 | 0.05 |
| FMM | 0.35 | 0.125 | 0.2 |
| MFM | 0.35 | 0.125 | 0.2 |
| MMF | 0.35 | 0.125 | 0.2 |
| FFM | 0.2 | 0.125 | 0.35 |
| FMF | 0.2 | 0.125 | 0.35 |
| MFF | 0.2 | 0.125 | 0.35 |
| FFF | 0.05 | 0.25 | 0.5 |

***Table S9:*** *Fitness of simulated offspring was based on the additive effects of their multi-locus haplotypes (fitness effect hapotype1 + fitness effect haplotype 2), and the habitat in which they live. This number was used to assign survival probabilities to each offspring.*

### Supplementary Methods

#### 1) Investigating the effect of heterochiasmy and recombination on the spread of adaptive mutations through hybrid zones using forward simulations.

We used population genetic simulations to investigate how heterochiasmy and recombination suppression influence the spread of freshwater adaptive alleles to neighbouring freshwater populations via hybrid zones. Specifically we used a five population model to simulate a selection-migration balance maintaining divergence among a freshwater (population 1), intertidal (population 2), and marine population (population 3). A second hybrid zone with intertidal (population 4) and freshwater population (population 5) was included with connection to the first via marine population (population 3), resulting in parallel hybrid zones with access to the same marine population.

```
       Population1 <-> Population2 <-> Population3 <-> Population4 <-> Population5
       (freshwater)    (intertidal)      (marine)      (intertidal)    (freshwater)
Ne:        200             200             600             200             200


chromosome homolog1:           Locus1 ----- Locus2 ----- Locus3
                                        X            X   sex-specific recombination rate
chromosome homolog2:           Locus1 ----- Locus2 ----- Locus3        (rr.female, rr.male)
```

Using diploid forward simulations to track alleles at three linked autosomal loci of equal recombination distance apart, we simulated for each of 120 generations:

1. the migration of individuals into and out of each population (m=0.2*Ne individuals per generation per year with a female-biased migration ration of 3females:1males mimicking ratios found in migrating marine fish);

2. for each individual, the production of haploid gametes via meiosis with sex-specific recombination rate among neighboring loci specified by probabilities (rr.male and rr.female)

3. the pairing of male and female gametes to produce 100 diploid offspring per breeding pair (clutch size = 100).

4. strong selection on offspring with survival dependent on the additive effects of the multilocus haplotypes carried by each individual and the habitat in which they are found (e.g. in freshwater and marine habitats selection favors the additive number of freshwater and marine alleles respectively; in intertidal populations selection favors individuals with cis-linked haplotypes of marine and/or freshwater alleles on both chromosome homologs; see Table S9 for more information)

5. finally to maintain equal population sizes, we imposed a population carrying capacity of 200 by randomly selecting 200 of the surviving offspring from all breeding pairs in the current generation (g) to continue as breeders in the subsequent generation (g+1).

We started each simulation with a two-locus secondary contact scenario where Population 1 carried freshwater adaptive alleles at Locus 1 and 3, while all other populations carried marine alleles. (Locus 2 is initially invariant among all populations, with no effect on offspring survival). We allowed migration, breeding (with recombination) and selection to occur for 30 generations during which a selection-migration balance is quickly established and maintains divergent allele frequencies in Population 1 and 3. Even with the maintenance of this divergence, the freshwater adaptive allele spreads from population 1 through the hybrid zone into population 3 and subsequently invades the parallel hybrid zone (populations 4 and 5) with a second selection-migration balance maintaining divergence in allele frequencies between populations 3 and 5. Then, after 30 generations, a new freshwater-beneficial mutation was introduced at Locus 2, by the arrival into freshwater Population 1 of N=0.01*ne migrants homozygous for freshwater alleles at all three loci. With locus 2 no longer neutral, from this point on the survival probabilities of offspring now depended on their three locus haplotypes.

We ran 8 different simulations that differed in the extent of heterochiasmy and recombination suppression in cis-(coupled)-heterozygotes (see Table S8), and for each simulation tracked allele frequencies at each locus in each population. Each of the 8 simulations was replicated 16 times. We were specifically interested in differences in how quickly the new freshwater adaptive mutation at Locus 2 was able to establish and increase to high frequency in Population 1 (where it was first introduced), as well as how the different parameters influenced the speed at which this new allele was able to spread and establish in Population 5 (the freshwater population in the neighbouring hybrid zone; see Sup Figs 6-13). Simulations were written and run in R.

Under all simulations, recombination suppression in cis(coupled)heterozygotes significantly increases the rate at which adaptive alleles establish in the population into which they are introduced (eg. compare simulation 1 and 2), and for the most part the rate at which they spread through hybrid zones into new freshwater populations (e.g compare simulation 7 and 8). In contrast, strong heterochiasmy compared to no heterochiasmy slows the rate at which adaptive alleles establish in a population (eg. 1 vs 3), but has little effect when there is recombination suppression in cis(coupled)heterozygotes (2 vs 4). When considering heterochiasmy as a derived recombination-suppression state in males, compared to an ancestral state where males recombine equally fast as females (simulation 3 vs 5), heterochiasmy is slightly more likely to facilitate the establishment of a new allele in the population into which it was introduced, as well as the spread of that allele to other populations. However if heterochiasmy is viewed as a derived state of increased recombination in females compared to an ancestral state of low recombination in both sexes (e.g. 3 vs 7), it significantly decreases the rate at which alleles are fixed but not the rate at which they spread.

#### 2) Home-made library preparation protocol

This protocol is optimized for high-throughput preparation of genomic DNA libraries compatible for sequencing on Illumina HiSeq 3000. All DNA size selections and cleanups were performed following Ampure XP® (Beckman-Coulter) SPRI bead size selection protocol.

**Step 1: DNA fragmentation using Covaris® LE220 instrument**

- Dilute 300-500 ng of good quality genomic DNA in 1X TE buffer to a total volume of 130 µl and transfer to covaries 96 well microtube plate (SKU:520078)

- Insert sample filled plate into the Covaris® LE220 plate holder and run with the following settings to obtain an average fragment size of 300bp.

| | |
|---|---|
| Sample volume | 130µl |
| Duty factor | 30% |
| Peak Incident Power | 450 |
| Cycles per Burst | 200 |
| Treatment time | 80 sec |

- Retrieve sheared samples from Covaris® plate into a 96 well plate and perform bead clean up. The following method is designed to yield fragments of size 300 bp or above.

| | |
|---|---|
| Input DNA: | 130µl |
| SPRI volume | 104µl (0.8 times the sample volume) |
| Resuspension volume | 30µl |
| Elution volume | 30µl |

**Step 2: End repair**

- Transfer 15µl of the eluate from above into a fresh plate for end repair
- Prepare the following reaction mix and add it to the DNA

| **Reagent** | **1x** |
|---|---|
| Water | 3.75 µl |
| 10X T4 DNA ligase buffer | 2.5 µl |
| 10mM dNTP mix | 1 µl |
| T4 DNA polymerase | 1.25 µl |
| Klenow DNA polymerase | 0.25 µl |
| T4 Polynucleotide kinase | 1.25 µl |

| | |
|---|---|
| **Total volume** | **10 µl** |

- Add 10 µl of reagent into 15 µl of sample. Mix well and spin down. Incubate at room temperature for 30 minutes
- Perform DNA size selection to remove fragments larger than 500 bp.

| | |
|---|---|
| Input DNA | 25 µl |
| SPRI volume | 15 µl (0.6 times the sample volume) |
| PEG to save | 40 µl (saving fragments < 500 bp) |
| SPRI volume | 15 µl (add to the saved PEG to extract all smaller sized DNA) |
| Resuspension volume | 18 µl |
| Final elution volume | 17 µl |

- Store samples in fridge if not proceeding right away

**Step 3: A-tailing**

- Prepare the following master mix and add it to the sample

| **Reagent** | **1x µl** |
|---|---|
| NEB Buffer 2 | 2.5 µl |
| 1 mM dATP | 0.5 µl |
| Klenow exo- | 0.5 µl |
| Water | 4.5 µl |
| **Total** | **8 µl** |

- Incubate at 37$^0$C for 30 minutes, then heat inactivate at 75$^0$C for 20 minutes
- Save 1 µl sample for running on bioanalyzer or 2 µl for checking on gel
- Directly proceed to the next step without waiting

**Step 4: Adapter ligation**

- Mix 1 µl 10 µM adapters with DNA sample (Illumina TruSeq single index adapter ordered from IDT)
  Note down the unique barcode used for each sample.
- Add the following ligase master mix to each sample

| **Reagent** | **1x** |
|---|---|
| 10mM rATP | 3 µl |
| T4 DNA ligase | 1.5 µl |
| Water | 0.5 µl |

| | |
|---|---|
| **Total** | **5 µl** |

- Incubate15 minutes at $20^0$C. Then 5 minutes at $65^0$C to inactivate the ligase. Allow samples to cool before proceeding

- Perform bead clean-up to remove the enzymes

| | |
|---|---|
| DNA volume: | 31 µl |
| SPRI volume | 24.8 µl (0.8X the sample volume) |
| Resuspension volume | 15 µl |
| Elution volume | 14 µl |

**Step 5: Library enrichment by PCR**

- Add the following ligase master mix to each sample

| **Reagent** | **1x** |
|---|---|
| Water | 2.75 µl |
| 5x Buffer | 5 µl |
| 10mM dNTP | 0.5 µl |
| 10µM Truseq PCR1 | 1.25 µl |
| 10µM Truseq PCR2 | 1.25 µl |
| Phusion polymerase | 0.25 µl |
| **Total** | **11 µl** |

- Perform PCR reaction with the following condition

| | | |
|---|---|---|
| $95^0$C | 30 sec | 1 cycle |
| $98^0$C | 15 sec | 6 cycles |
| $62^0$C | 30 sec | |
| $72^0$C | 30 sec | |
| $72^0$C | 10 min | 1 cycle |
| $4^0$C | Hold | Infinitely |

**Step 6: Library validation and pooling**

- Confirm adapter ligation by running the sample on bio-analyzer or checking on gel (load 2 µl sample). Successful adapter ligation will increase the fragment size by ~120 bp.
- Measure final library concentration by TECAN plate reader (using picogreen dye)
- Pool equal quantity of all libraries to be sequenced in a lane

- Perform bead clean up on pooled libraries to remove PCR reagents

  If pooled sample volume is less than 50 µl, make up to 50 µl by adding EB buffer

| | |
|---|---|
| Sample volume | 50 µl |
| SPRI volume | 40 µl (0.8 times the sample volume) |
| Resuspension volume | 61 µl |
| Elution volume | 60 µl |

- Perform DNA double size selection to remove fragments larger than 600 bp and smaller than 300 bp

| | |
|---|---|
| Input DNA | 60 µl |
| SPRI volume | 24 µl (0.4 times the sample volume) |
| PEG to save | 84 µl (saving fragments < 600 bp) |
| SPRI volume: | 18 µl (add to the saved PEG to select all fragments >300 bp) |
| Resuspension volume | 31 µl |
| Final elution volume | 30 µl |

- Check the quality of the library pools by bio-analyzer (example shown below) and measure the quantity by qubit high-sensitivity reagent.

- Submit 2.5nM library for sequencing

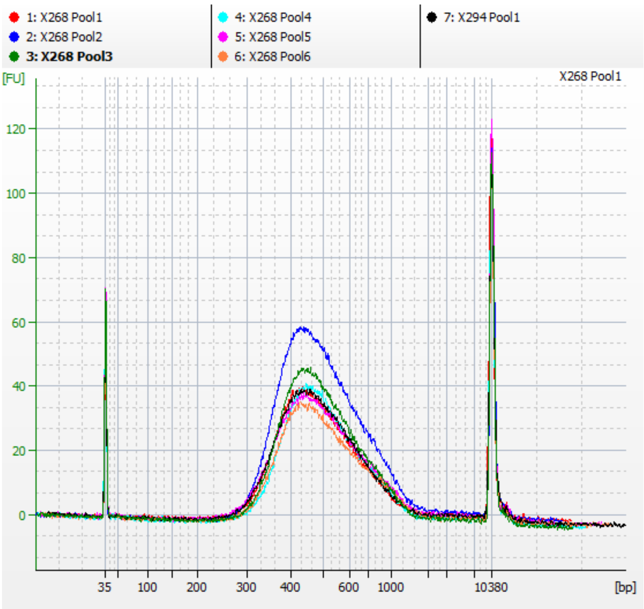


*Bioanalyzer profile of size-selected library pools. Library size profiles of seven pools are shown. All of them have an average fragment size of about 420 bp (300 bp insert + 120 bp adapter).*

Reagents used for library preparation:

| **Reagent** | **NEB catalogue number** |
|---|---|
| T4 DNA polymerase | M0203L |
| T4 Polynucleotide kinase | M0201L |
| Klenow exo- | M0212L |
| T4 DNA ligase | M0202L |
| Klenow DNA polymerase | M0210L or M0210S |
| Phusion High-Fidelity DNA polymerase | M0530L |
